# The UTX Tumor Suppressor Directly Senses Oxygen to Control Chromatin and Cell Fate

**DOI:** 10.1101/510479

**Authors:** Abhishek A. Chakraborty, Tuomas Laukka, Matti Myllykoski, Alison E. Ringel, Matthew A. Booker, Michael Y. Tolstorukov, Yuzhong Jeff Meng, Sam Meier, Rebecca B. Jennings, Amanda L. Creech, Zachary T. Herbert, Jessica Spinelli, Samuel K. McBrayer, Benjamin A. Olenchock, Jacob D. Jaffe, Marcia C. Haigis, Rameen Beroukhim, Sabina Signoretti, Peppi Koivunen, William G. Kaelin

**Affiliations:** Department of Medical Oncology, Dana-Farber Cancer Institute and Brigham and Women’s Hospital, Harvard Medical School, Boston, MA 02215, USA.; Biocenter Oulu, Faculty of Biochemistry and Molecular Medicine, Oulu Center for Cell-Matrix Research, University of Oulu, FIN-90014 Oulu, Finland.; Department of Cell Biology, Harvard Medical School, Boston, MA 02115, USA.; Department of Informatics, Dana-Farber Cancer Institute, Harvard Medical School, Boston, MA 02215, USA.; Broad Institute of Harvard and MIT, Cambridge, MA 02142, USA.; Biological and Biomedical Sciences, Harvard Medical School, Boston, MA 02115, USA.; Department of Cancer Biology, Dana-Farber Cancer Institute, Boston, MA, Harvard Medical School, Boston, MA 02115, USA.; The Harvard-MIT Program in Health Sciences and Technology, Harvard Medical School, Boston, MA.; Department of Pathology, Brigham and Women’s Hospital, Harvard Medical School, Boston, MA 02115, USA.; Molecular Biology Core Facility, Dana-Farber Cancer Institute and Brigham and Women’s Hospital, Harvard Medical School, Boston, MA 02215, USA.; Division of Cardiovascular Medicine, Department of Medicine, The Brigham and Women’s Hospital, Boston, MA 02115, USA; Harvard Medical School, Boston, MA 02115, USA.; Howard Hughes Medical Institute, Chevy Chase, MD 20815, USA.

## Abstract

Mammalian cells express multiple 2-oxoglutarate (OG)-dependent dioxygenases, including many chromatin regulators. The oxygen affinities, and hence oxygen sensing capabilities, of the 2-oxoglutarate (OG)-dependent dioxygenases reported to date vary widely. Hypoxia can affect chromatin, but whether this reflects a direct effect on chromatin-modifying dioxygenases, or indirect effects caused by the hypoxic-induction of the HIF transcription factor or the endogenous 2-OG competitor 2-hydroxyglutarate (2-HG), is unclear. Here we report that hypoxia induces a HIF- and 2-HG-independent histone modification signature consistent with KDM inactivation. We also show that the H3K27 histone demethylase KDM6A (also called UTX), but not its paralog KDM6B, is oxygen-sensitive. KDM6A loss, like hypoxia, prevented H3K27me3 erasure and blocked differentiation. Conversely, restoring H3K27me3 homeostasis in hypoxic cells reversed these effects. Therefore, oxygen directly affects chromatin regulators to control cell fate.

**One Sentence Summary:** KDM6A demethylase activity is diminished under hypoxic conditions and causes changes in gene expression programs that govern cell fate.

---

The appearance of oxygen in earth’s atmosphere was a terrestrial milestone and created the evolutionary selection pressure for a conserved pathway used by metazoans to sense and respond to changes in ambient oxygen. Central to this pathway is a heterodimeric transcription factor called Hypoxia-Inducible Factor (HIF), which consists of an unstable α-subunit and a stable β-subunit (HIF1β or ARNT) (*1-3*). Under well-oxygenated conditions, the α-subunit is prolyl hydroxylated by members of the EglN family of 2-oxoglutarate (2-OG)-dependent dioxygenases. Hydroxylated HIFα is then polyubiquitylated by the pVHL tumor suppressor protein and subsequently degraded. Hypoxia inactivates the EglNs and thereby stabilizes HIFα, which then associates with ARNT and transcriptionally activates genes that promote adaptation to inadequate oxygen (*1-3*).

The 2-OG-dependent dioxygenase family also includes the collagen prolyl hydroxylases, the JmjC domain histone demethylases (KDMs), the TET DNA hydroxylases, and ~50 other enzymes that are relatively understudied (*4*). In contrast to the high oxygen affinity collagen prolyl hydroxylases (*5-7*), the EglNs exhibit relatively low oxygen affinities (high *K_M_*) (*8-11*), which enables them to sense physiological changes in oxygen. The oxygen affinities of most of the other family members are unknown.

Many KDM genes are transcriptionally activated by hypoxia and, where studied HIF, perhaps to compensate for a decrease in their specific activity (*12-18*). In support of the latter, some KDMs have been reported to have low oxygen affinities in vitro (*19-21*). Finally, hypoxia promotes the accumulation of hypermethylated histones (*20-28*).

These considerations suggest that oxygen directly regulates KDM function. A caveat, however, is that hypoxia (and HIF) could also indirectly affect KDM activity. For example, in some systems hypoxia causes the accumulation of the L-enantiomer of 2-hydroxyglutarate (L-2-HG) (*29-32),* which is an endogenous inhibitor of 2-OG-dependent dioxygenases. Secondly, HIF transcriptionally regulates many chromatin modifying enzymes (*12-18*). Thirdly, HIF can induce profound changes in cell state that might indirectly affect chromatin, such as by upregulating transcription factors that enforce an epithelial-mesenchymal transition (EMT) (*33*). It therefore remains unclear whether the changes in histone methylation observed in hypoxic cells in vivo reflect a direct effect of hypoxia on KDMs or are an indirect consequence of hypoxia.

To address this question, we lentivirally transduced an *Arnt*-defective (and thus HIF-inactive) mouse hepatoma cell line (mHepa-1 c4) (*34*), to express either Green Fluorescent Protein (GFP), wild-type ARNT (WT), or a functionally inactive (and possibly less stable) ARNT mutant (Δ414) that is missing 414 base pairs from its N-terminus, thereby eliminating its basic-Helix-Loop-Helix (bHLH) domain (Fig. 1, A and B, and fig. S1A). mHepa-1 c4 did not tolerate long term growth in 1% oxygen, which is the oxygen concentration typically used to model hypoxia *ex vivo.* We therefore used more modest levels of hypoxia (2-5%) to study these cells. Importantly, we confirmed that wild-type ARNT, but not ARNT Δ414, coimmunoprecipitated with endogenous HIF1α (fig. S1B), and that canonical HIF-target genes (e.g. *Egln3* and *Ndrg1*) were transcriptionally induced by 5% oxygen in the mHepa-1 c4 cells expressing wild-type ARNT, but not in the cells expressing ARNT (Δ 414) or GFP (Fig. 1C and fig. S1C).

**Fig. 1.**
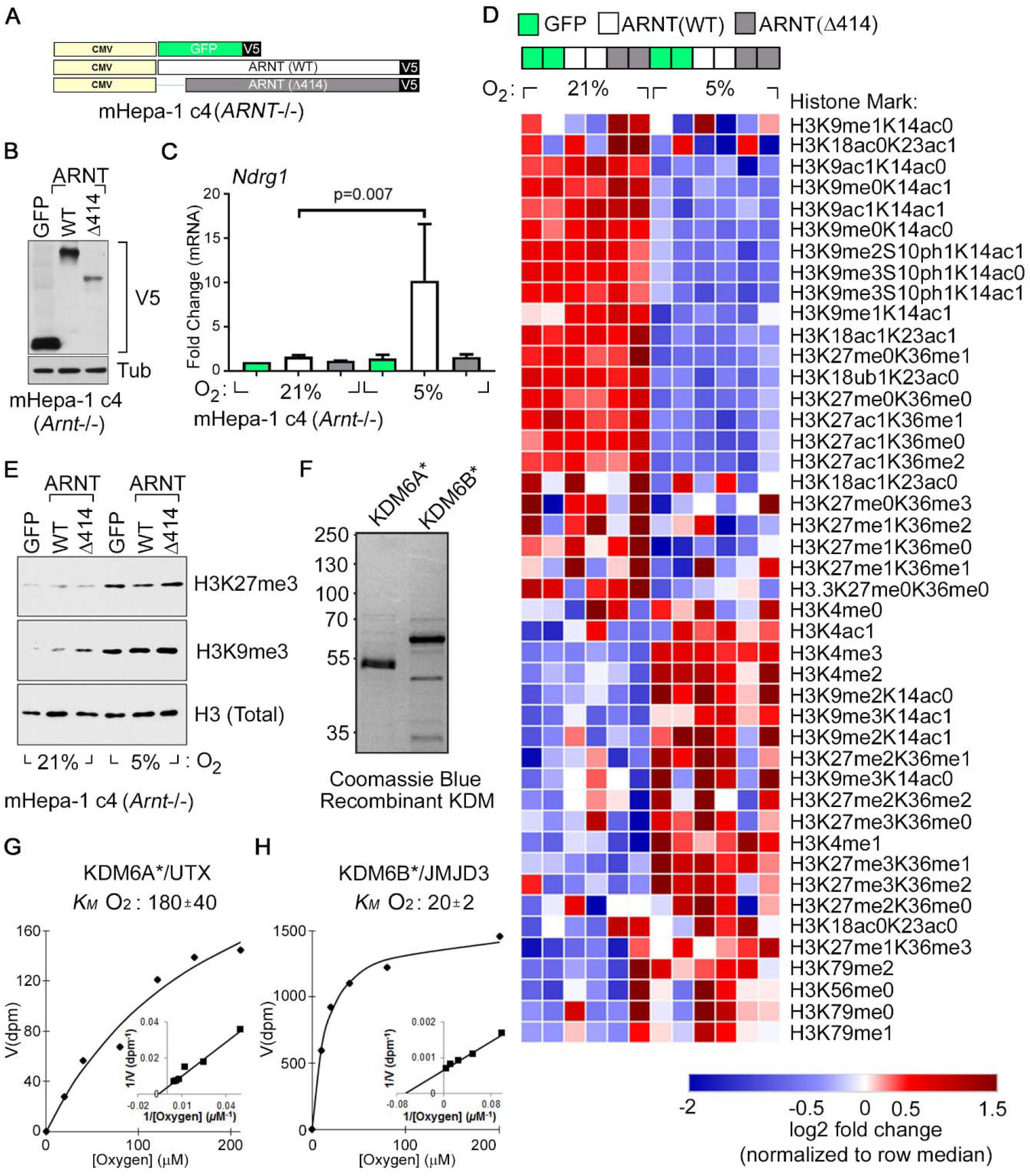
Hypoxia Causes HIF-Independent Histone Hypermethylation. (**A-E**) Vector schematic (**A**), Immunoblot analysis (**B**), Real-Time qPCR analysis (**C**), Histone modification profiling by mass spectrometry (**D**), and Histone immunoblot analysis (**E**) of *Arnt-* deficient mouse Hepatoma (mHepa-1 c4) cells that were lentivirally transduced to produce the indicated V5-tagged proteins and cultured at the indicated oxygen levels for 1 day (**C**) or 4 days (**D** and **E**). In (**B**), Tub = Tubulin. In (**C**), data represent mean±SD (n=3) and p-value was calculated using the Students *t*-test. In (**D**), columns represent two biological replicates of the indicated samples and the color in each cell represents log2 fold change relative to all samples in the row, normalized for total histone using an internal control peptide (Histone H3: residues 41-49). (**F**-**H**) Coomassie blue staining (**F**) and biochemical analysis of baculovirally expressed and purified JumonjiC (JmjC) catalytic domains of KDM6A [KDM6A*] (**G**) and KDM6B [KDM6B*] (**H**). *K_M_* values are mean±SD (n=3 independent measurements).

We next used a previously characterized multiplexed mass spectrometric platform to monitor changes in histone methylation in response to hypoxia in the isogenic mHepa-1 c4 cells in which HIF function was or was not restored. This assay simultaneously quantifies many histone modifications by comparing their abundance on tryptic digests of Histone H3 to spiked-in internal control peptides harboring the corresponding modification (*35*).

Unsupervised clustering of histone modification patterns revealed that the isogenic cell lines clustered primarily based on oxygen availability during growth and not HIF status (Fig. 1D). Consistent with prior reports (*20-28*), hypoxia promoted the hypermethylation (me2/me3 states) of H3K4, H3K9, and H3K27 (Fig. 1D). Hypermethylation of H3K9 and H3K27 was also confirmed by immunoblot analysis (Fig. 1E). We also observed a concomitant decrease in hypomethylated H3K27 (me0/me1 states) and acetylated H3K27, in response to hypoxia (Fig. 1D). Hypermethylation and decreased acetylation are both consistent with decreased histone demethylase activity because histone methylation and acetylation are reciprocally regulated. The changes in H3K27 methylation were not explained by obvious increases in the expression of EZH2, which controls bulk H3K27 methylation, or decreased expression of the primary H3K27 histone demethylases KDM6A and KDM6B (fig. S2).

Hypoxia also promoted histone hypermethylation in *VHL^−/−^* RCC4 cells, which constitutively contain high HIF levels (fig. S3). Thus hypoxia promotes histone hypermethylation both in cells that cannot mount a HIF response (mHepa-1 c4 cells) and in cells that constitutively overproduce HIF (RCC4 cells), arguing that the effects of hypoxia on histone methylation are not caused by changes in HIF activity.

In some, but not all, cell types hypoxia induces the accumulation of L-2-HG (*6, 29-32*), which is a potent inhibitor of 2-OG-dependent dioxygenases. We therefore asked if the effects of hypoxia on histone methylation in our model were caused by a HIF-independent induction of L-2-HG or perhaps an alternative metabolic change, such as decreased 2-OG, which also can affect KDM activity (*36, 37*). In the isogenic mHepa-1 c4 cells, exposure to 5% oxygen did not significantly induce either total 2-HG, measured absolutely and relative to succinate (fig. S4A), or enantiomer-specific 2-HG (fig. S4, B and C), and, if anything, increased 2-OG levels (fig. S4D). 2-HG was modestly induced in parental mHepa-1 c4 cells by more profound levels of hypoxia (0.5-2% oxygen) (fig. S4, C and E), albeit as D-2HG rather than L-2HG (fig. S4C). The significance of this latter finding is unclear. Even under these more extreme conditions the 2-HG levels achieved were ~100-fold below both the intracellular levels in mutant IDH1 cells (fig. S4F) and the intracellular levels required to promote histone methylation in mHepa-1 c4 cells treated with esterified versions of D-2HG or L-2HG (fig. S4, G and H). The latter observation is consistent with the biochemical D-2-HG and L-2HG IC_50_ values for the KDMs reported to date (*38*).

Hypoxia can induce the production of reactive oxygen species (ROS), which can also inhibit 2-OG-dependent dioxygenase function (*39*). In dose titration experiments with the ROS-inducer *tert*-Butyl Hydroperoxide (tBHP), we found that intracellular ROS levels ~10-fold higher than observed with 2% oxygen were required to induce histone methylation (fig. S5). Together, these findings suggested that the HIF-independent effects of hypoxia on KDM activity were not caused by increased L-2-HG, decreased 2-OG, or increased ROS, but instead were caused by a direct effect of hypoxia on the enzymatic activities of specific KDMs. In support of this idea, we found that recombinant KDM4B, KDM5A, and KDM6A have relatively low oxygen affinities (high oxygen *K_M_* values) that are comparable to the EglN family, while recombinant KDM4A, KDM5B, KDM5C, KDM5D, and KDM6B have high oxygen affinities (low oxygen *K_M_* values) (Fig. 1, F to H, and fig. S6 to S8). We had difficulty purifying stable full-length KDM6B, but we confirmed that both full-length KDM6A and the isolated KDM6A catalytic domain had low oxygen affinities compared to their KDM6B counterparts (Fig. 1, F to H and fig. S8).

We focused our attention on the KDM6 H3K27 demethylases because both hypoxia and H3K27 methylation have been linked to the control of stem cell biology and differentiation (*40, 41*) and because of the availability of pharmacological tools for manipulating this histone mark. To begin addressing the physiological relevance of our findings, we first confirmed that hypoxia induced H3K27 methylation in multiple additional cell lines including 293T embryonic kidney cells, MCF7 breast cells, and SK-N-BE(2) neuroblastoma cells (fig. S9). Moreover, histologic analysis showed elevated H3K27 methylation in mouse tissues that are known to be hypoxic, such as the kidney (*42*), splenic germinal centers (*43*), and thymus (*44*), but not in well-oxygenated tissues such as in the heart (Fig. 2A). Similarly, and in keeping with prior studies (*24, 27, 28*), H3K27 methylation was increased in hypoxic regions of mouse tumors (fig. S10). Finally, Gene-Set Enrichment Analysis (*45*) of gene expression data from ~2000 human tumors that were previously annotated as “Normoxic” or “Hypoxic” based on their HIF signature (*6*) (Table S1) revealed that “Hypoxic” tumors had transcriptional signatures indicative of H3K27 hypermethylation (Fig. 2, B to D, and Tables. S2 and S3).

**Fig. 2.**
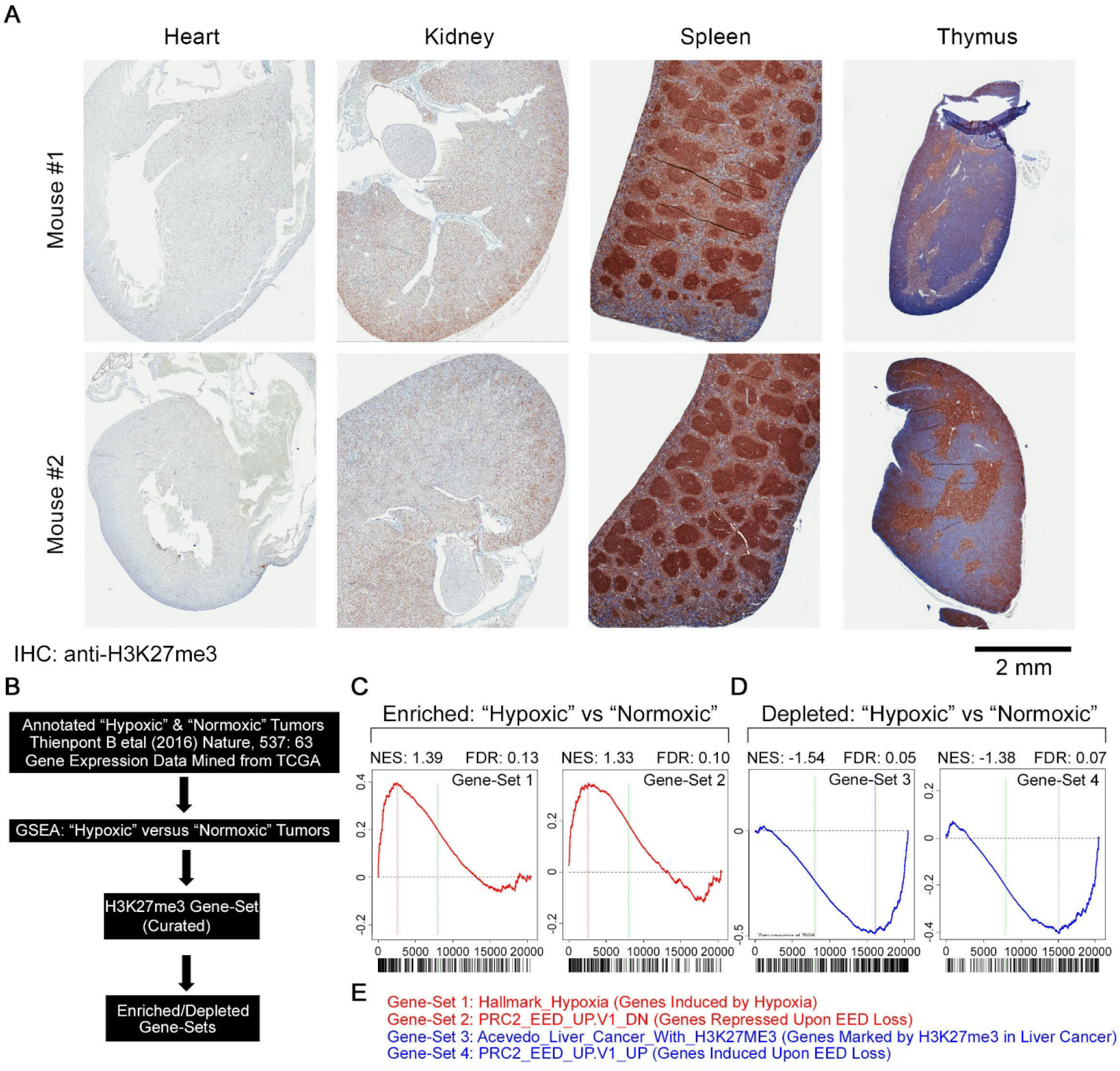
Physiological Hypoxia Promotes H3K27 Hypermethylation. (**A**) Immunohistochemical analysis of the indicated tissues derived from representative male (*76*) and female (*77*) age-matched mice. (**B-D**) Analytical flowchart (**B**) and Gene Set Enrichment Analysis showing the enrichment plots for transcriptional signatures that are enriched (**C**) or depleted (**D**) in “hypoxic” tumors. (**E**) Description of gene sets shown in (**C**) and (**D**).

In many models, such as the well-studied C2C12 myoblast differentiation model wherein cells are shifted from growth media (GM) to differentiation media (DM), hypoxia blocks differentiation. In the C2C12 model differentiation can be easily scored based on the appearance of Myosin Heavy Chain (MyHC) positive multinucleated myotubes. We confirmed earlier reports (*46, 47*) that hypoxia blocks C2C12 differentiation (Fig. 3, A and B, and fig. S11, A and B). The hypoxic C2C12 cells grown in DM entered a quiescence-like state but more readily proliferated when returned to GM under normoxic conditions compared to cells that had been induced to differentiate under normoxia (fig. S11, C-E). Similarly, hypoxia blocked the myogenic differentiation of mouse embryo fibroblasts engineered to conditionally express MyoD (fig. S12).

**Fig. 3.**
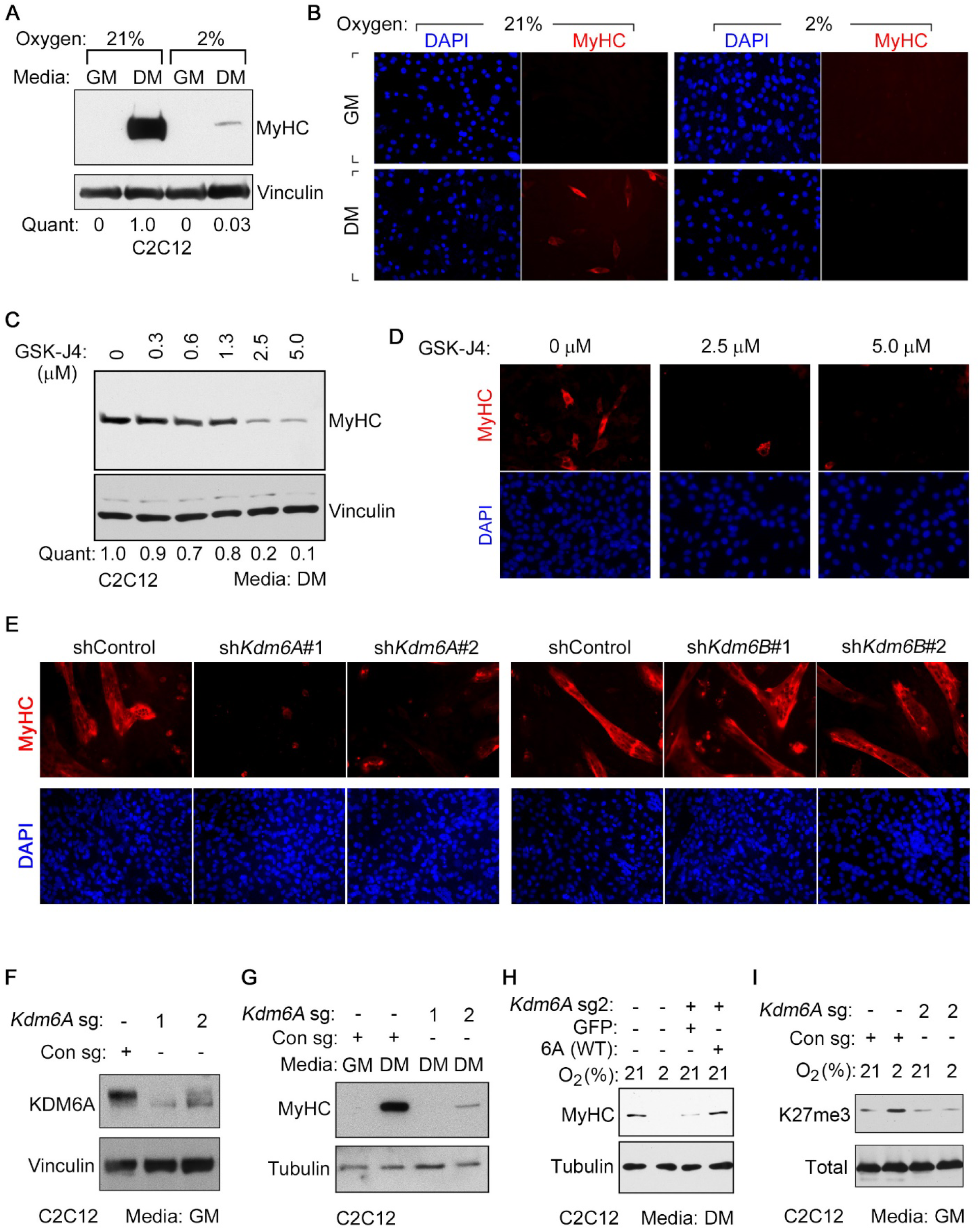
KDM6A Loss Mimics the Myogenic Differentiation Block Caused by Hypoxia. (*58*) Immunoblot (**A**) and Immunofluorescence analysis (**B**) of mouse C2C12 myoblasts cultured in either growth media (GM) or differentiation media (DM) for 4 days at the indicated oxygen concentrations. MyHC = Myosin Heavy Chain. (**C-D**) Immunoblot (**C**) and Immunofluorescence analysis (**D**) of C2C12 cells cultured in DM for 4 days in the presence of the indicated concentrations of GSK-J4. (**E**) Immunofluorescence analysis of C2C12 cells that were lentivirally transduced to express the indicated shRNAs and then cultured in DM for 4 days. (**F-G**) Immunoblot analysis of C2C12 cells lentivirally transduced to express the indicated sgRNAs and cultured either in GM (**F**) or in the indicated media (**G**) for 4 days. (**H**) Immunoblot analysis of C2C12 cells expressing, where indicated, *Kdm6a* sg2 [described in (**F**) and (**G**)] that were lentivirally transduced to produce either GFP (control) or wild-type human KDM6A [6A(WT)] and then cultured in DM at the indicated oxygen concentrations for 4 days. The mouse *Kdm6A* sg2 target sequence is not conserved in human *KDM6A.* (**I**) Histone immunoblot analysis of C2C12 cells expressing the indicated sgRNA that were cultured at the indicated oxygen concentrations for 3 days. In (**A**) and (**C**), “Quant” represents fold-change in densitometric ratios of MyHC (normalized to Vinculin) relative to untreated normoxic cells cultured in DM.

Eliminating ARNT in C2C12 cells using CRISPR/Cas9 blocked (rather than accentuated) their ability to differentiate under normoxic conditions (fig. S13, A to C), consistent with an earlier study that used a HIF1α siRNA (*48*), and did not rescue their ability to differentiate under hypoxia (fig. S13D). Moreover, expressing stabilized versions of HIF1α or HIF2α (*49*) did not block normoxic C2C12 differentiation (fig. S13, E and F). Therefore the differentiation block exhibited by hypoxia C2C12 cells is not due to HIF activation.

Total 2-HG was not induced, and L-2HG was induced only about 2-fold (fig. S14, A to C), in C2C12 cells by 2% oxygen. C2C12 with intracellular L-2HG levels that were 3-5-fold higher than observed under hypoxia, achieved using esterified L-2HG or elimination of the L-2HG catabolic enzyme L2HGDH using CRISPR/Cas9, still differentiated (fig. S14, D to G). Therefore, the effects of hypoxia on C2C12 differentiation were not caused by 2-HG.

KDM6A and KDM6B are the primary H3K27 demethylases in cells (*50*). Similar to hypoxia, treating C2C12 cells with GSK-J4, a dual pharmacological inhibitor of KDM6A and KDM6B, promoted H3K27 hypermethylation and blocked myogenic differentiation (Fig. 3,C and D, and fig. S15). This was specific because treating C2C12 cells with KDM-C70, a pan-inhibitor of the KDM5 family, did not block differentiation (fig. S16). Similar results were obtained with MEFs expressing MyoD (fig. S12).

The differential oxygen affinities of KDM6A and KDM6B suggested that the ability of hypoxia to promote H3K27 methylation and block differentiation is caused specifically by a loss of KDM6A activity. To specifically address the contribution of KDM6A we identified effective shRNAs against *Kdm6a* and *Kdm6b*. Importantly, downregulation of KDM6A with two different shRNAs, but not downregulation of KDM6B, phenocopied the effects of hypoxia on differentiation (Fig. 3E and fig. S17, A to D), consistent with an earlier study (*51*). Moreover, eliminating KDM6A in C2C12 cells with CRISPR/Cas9 blocked their ability to differentiate unless they were rescued with an sgRNA-resistant KDM6A cDNA (Fig. 3, F to H). This was specific because eliminating KDM6B had no effect (fig. S17, E and F) and eliminating KDM5A actually promoted differentiation (fig. S18). Notably, bulk H3K27 trimethylation was insensitive to changes in oxygen in the *Kdm6a*-deficient C2C12 cells, consistent with KDM6A being the primary oxygen sensor amongst the KDM6 paralogs (Fig. 3I).

Previous work showed that KDM6A is directly recruited to myogenic targets during differentiation (*51*). Therefore, we asked if differentiation programs driven by KDM6A activity involve transcriptional changes that depend on H3K27me3 elimination. Comparing transcriptional signatures of normoxic C2C12 cells that were cultured in GM to DM revealed profound differences in transcriptional output, particularly of muscle-specific target genes, resulting in a poor correlation between the transcriptional profiles obtained under these two media conditions (Fig. 4, A to C, and Table S5). Hypoxia (and the consequent differentiation block), however, blunted the transcriptional differences between these two conditions (Fig. 4A), which was associated with a failure of these cells to induce muscle-specific markers in DM (Fig. 4, B and C). H3K27me3 status typically represses transcription. The inability of C2C12 cells grown in DM to activate late myogenic genes (e.g.: *Actc, Myl1,* and *Myog*) under hypoxia correlated with a failure to erase H3K27me3 at those loci (Fig 4, D to F and fig. S19), presumably due to inactivation of KDM6A. In contrast, H3K4me3 actually decreased at late myogenic genes under hypoxia (fig. S19).

**Fig. 4.**
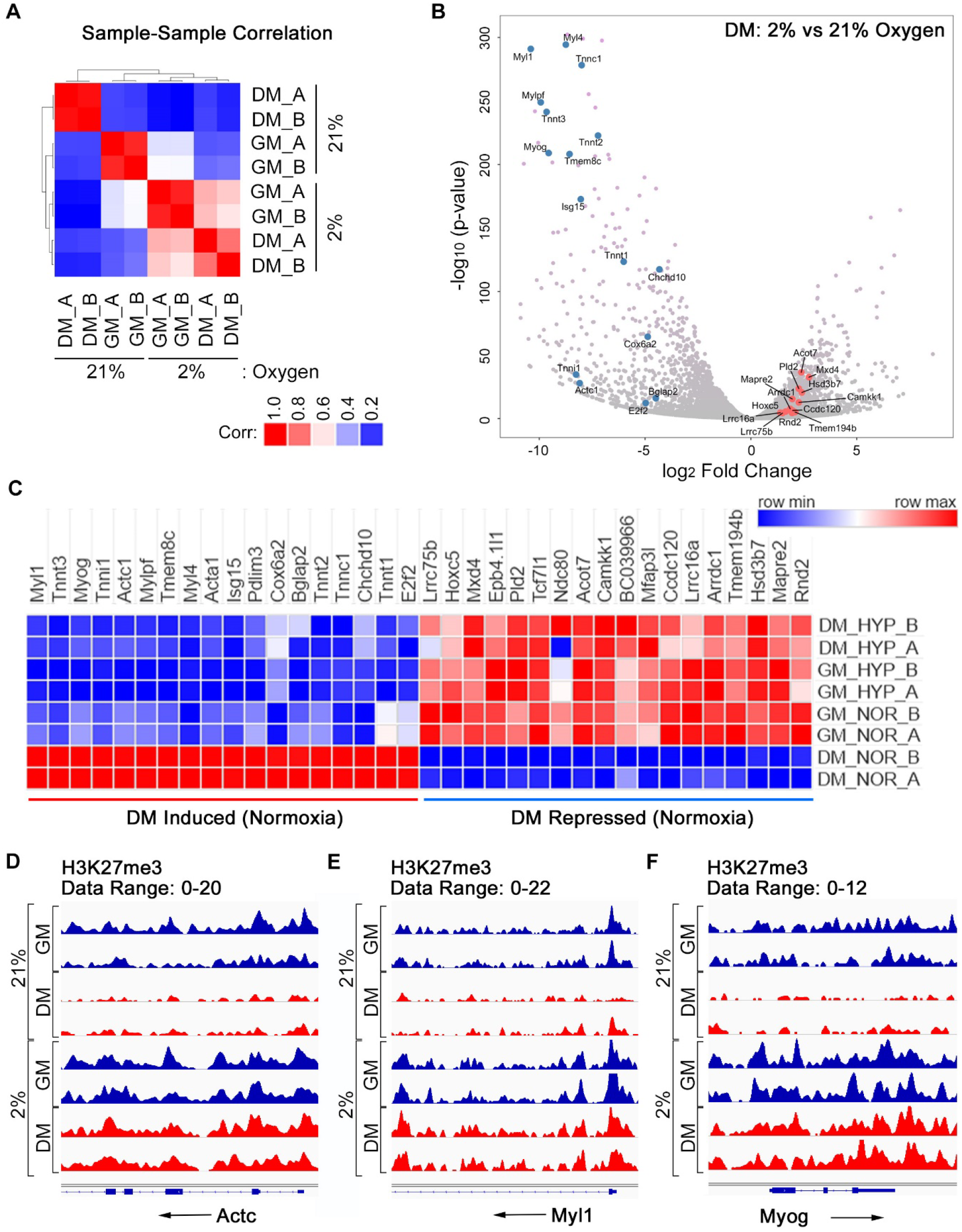
Hypoxia Blocks the Erasure of H3K27me3 and Induction of mRNAs Linked to Myogenic Differentiation. (**A**) Pearson correlation coefficient calculated by comparing the RNA-Seq transcriptional profiles obtained from two biological replicates (A and B) of C2C12 cells cultured in growth media (GM) or differentiation media (DM) for 4 days at the indicated oxygen concentration. (**B-C**) Volcano plot (**B**) and heatmap for selected mRNAs (**C**) showing differences in mRNA abundance, as determined by RNA-Seq, in C2C12 cells treated as in (**A**). (**D-E**) H3K27me3 levels at the *Actc1* (**D**), *Myl1* (**E**), and the *Myog* (**F**), locus as determined by ChIP-Seq analysis from C2C12 cells treated as in (**A**).

Loss of differentiation is a hallmark of cancer and *KDM6A* is a human tumor suppressor gene that is mutationally inactivated in a variety of cancers, including leukemias, kidney cancers, and bladder cancer (*52*). Remarkably, Gene Set Enrichment Analysis showed that a myogenic differentiation gene set, which presumably also contains genes linked to differentiation in other contexts, is more highly expressed in *KDM6*A wild-type bladder cancers compared to *KDM6A* mutant tumors (fig. S20 and Table S6).

These data suggest that KDM6A inactivation by hypoxia promotes the persistence of H3K27me3 and prevents the transcriptional reprogramming required for differentiation. If true, the effects of hypoxia on differentiation should be redressed by inhibiting H3K27 methyltransferase activity (Fig. 5A). Of note, and in keeping with our results using mHepa-1 c4 cells, hypoxia did not alter the protein levels of the EZH H3K27 methyltransferases in C2C12 cells (fig. S21A). Inhibiting the H3K27 methyltransferase EZH2 with either of two effective sgRNAs or with the pharmacological inhibitor GSK126 reduced H3K27me3 levels and partially rescued the ability of C2C12 cells to differentiate under hypoxic conditions (Fig. 5, B and C, and fig. S21, B and C). The latter was specific because the G9a/GLP methyltransferase inhibitor UNC638 was ineffective in this regard (Fig. 5C and fig. S21C). Finally, GSK126 rescued the hypoxia-induced differentiation block in human primary myoblasts and in MEFs expressing MyoD-ER (fig. S21, D and E).

**Fig. 5.**
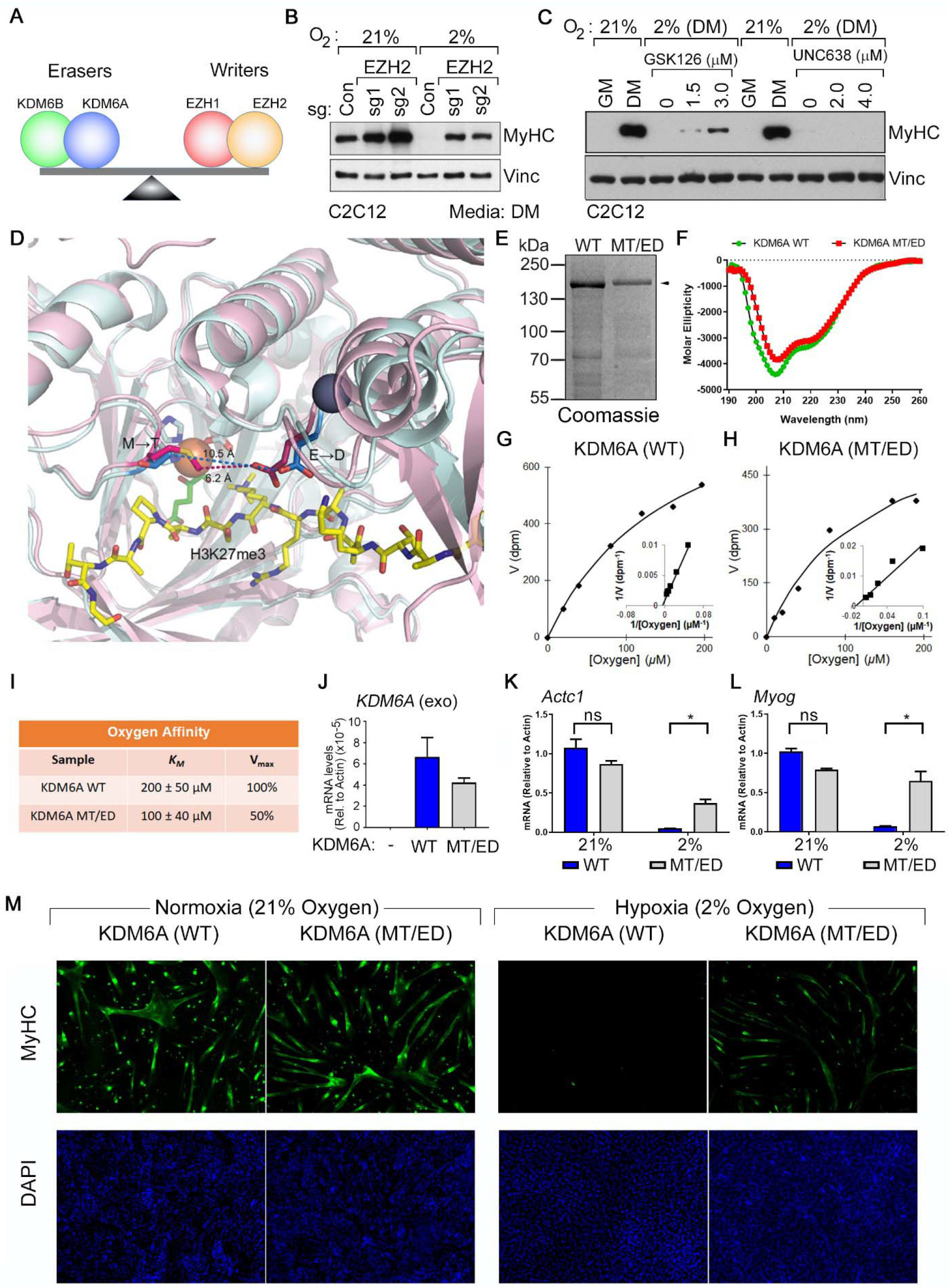
Restoring the Balance of H3K27 Methyltransferase Activity to H3K27 Demethylase Activity Rescues Myogenic Differentiation Under Hypoxic Conditions. (**A**) Model for control of H3K27 methylation by the indicated opposing demethylases (“erasers”) and methyltransferases (“writers”). (**B** and **C**) Immunoblot analysis of C2C12 cells lentivirally transduced to express the indicated sgRNAs (**B**)z or treated with the indicated histone demethylase inhibitors (**C**) during growth under the indicated conditions. (**D**) Structural models of the KDM6A (pink) and KDM6B (cyan) catalytic pockets. The non-conserved M^1190^ (KDM6A) T^1434^ (KDM6B) and the E^1335^ (KDM6A) D^1579^ (KDM6B) are highlighted. Peptidic H3K27me3 substrate (yellow), Fe^+2^ (orange), 2-oxo-glutarate (green), and Zn+^2^ (grey) are shown. (**E – I**) Coomassie Blue stained gel (**E**), Circular Dichroism analysis (**F**), Michaelis Menten plots (inset Lineweaver-Burke plot) (**G** and **H**), and measured oxygen *K_M_* values (**I**) of KDM6A wild-type and MT/ED double mutant. (**J-M**) mRNA levels of the indicated genes, relative to Actin, measured by Real-Time qPCR (**J to L**) and immunofluorescence analysis (**M**) of C2C12 cells transduced to produce wild-type human KDM6A or the KDM6A MT/ED variant and then cultured in DM at the indicated oxygen concentrations for 4 days.

Hypoxia can also, in at least a partially HIF-independent manner, alter the differentiation of human mammary epithelial (HMLE) cells (*24*), causing an EMT (fig. S22, A and B) and upregulation of the cancer stem-like marker CD44 (S22C). Similar to our findings in C2C12 cells, these hypoxia-associated changes were phenocopied by pharmacological (GSK-J4) (fig. S22, A to C) or by genetic (CRISPR/Cas9) disruption of KDM6A (fig. S22, D to F) and rescued by the EZH inhibitor GSK-126 (fig. S22, A to C).

To probe the link between oxygen availability, KDM6A activity, and differentiation further, we next treated C2C12 cells with the complex 1 inhibitor metformin. Although metformin is a pleotropic agent, we confirmed that it lowered oxygen consumption in hypoxic C2C12 cells (fig. S23A), which should increase intracellular oxygen availability. Metformin rescued the ability of C2C12 to lower H3K2me3 and to differentiate under hypoxia (fig. S23, B and C). This effect of metformin was specific because metformin did not rescue differentiation in C2C12 cells that lacked KDM6A (fig. S23D) and inhibited, rather than promoted, differentiation under normoxic conditions (fig. S23E).

Finally, we tried to directly increase KDM6A’s intrinsic oxygen affinity. We reasoned that certain non-conserved residues within their catalytic domains were responsible for the vastly different oxygen affinities of KDM6A and KDM6B. To identify these residues we overlaid previously published models of the catalytic JmjC domains of KDM6A with KDM6B (*53, 54*) and noted two non-conserved residues, M^1190^ (KDM6A) → T^1434^ (KDM6B) and E^1335^ (KDM6A) → D^1579^ (KDM6B), lining the 2-OG and Fe^+2^ binding pocket (Fig. 5D). As predicted, a KDM6A variant that harbored these two “KDM6B-like” changes [MT/ED: M^1190^ → T and E^1335^ → D], displayed a 2-fold increased affinity for oxygen in vitro, albeit at the cost of a decreased V_max_ (Fig. 5, E to I). Wild-type KDM6A and the MT/ED variant were comparably insensitive to L-2HG and to ROS, which was induced less than 2-fold by 2% oxygen in C2C12 cells (fig. S24, A-C). Reintroduction of either the KDM6A double mutant or wild-type KDM6A into KDM6A-deficient C2C12 cells rescued their ability to differentiate under normoxia (fig. S25A). The double mutant, however, was clearly superior to wild-type KDM6A at rescuing differentiation under hypoxic conditions in both parental and KDM6A-deficient C2C12 cells, presumably due to its enhanced oxygen affinity (Fig. 5, J to M, and fig. S25, B and C).

The importance of both oxygen and H3K27 methylation in regulating diverse biological processes, including embryological development, cellular differentiation, stemness, and malignant transformation, has been well described (*40, 55, 56*). Our findings argue that these observations are linked. Specifically, we argue that oxygen has both direct and indirect effects on chromatin and that the former involves enzymes such as KDM6A, which couple changes in oxygen availability to changes in H3K27 methylation, and ultimately affect the transcriptional control of cell fate. It is tempting to speculate, given the accumulation of H3K4 methylation and H3K9 methylation that we observed in hypoxic *Arnt*-deficient mHepa-1 c4 mouse hepatoma cells, together with our biochemical studies (and those of others), that at least one H3K4 and one H3K9 histone demethylase also act as oxygen sensors and contribute to chromatin regulation by oxygen. For example, our biochemical data, together with the data in the accompanying manuscript by Sonia Rocha, argue that inhibition of KDM5A contributes to the transcriptional response to hypoxia. Profound hypoxia can also increase DNA methylation in a HIF- and 2-HG-independent manner by inhibiting TET enzymes (*6*).

Our findings have potential implications for the importance of microenvironmental hypoxia in embryological development and stem cell maintenance as well as for the selection pressure to inactivate KDM6A in cancer. They also suggest that some of the biological effects of metformin, including its potential therapeutic effects for diseases such as cancer, are mediated at least partly by changes in KDM activity. In addition, hypoxia-induced epigenetic changes might contribute to the phenomenon of ischemic preconditioning, wherein a prior bout of ischemia confers protection against subsequent ischemic insults (*57*).

It is well established that the hypoxia can affect stemness and cellular differentiation by activating HIF and downstream target genes such as Oct4 (*55, 58*). Such effects are not mutually exclusive with a direct effect of oxygen on histone methylation and might, in fact, serve to reinforce on another. The HIF pathway is conserved throughout metazoan evolution but is not present in plants, yet hypoxia promotes stemness in both metazoans and plants (*59, 60*). It is possible that the oxygen sensing by histone demethylases evolutionarily preceded the emergence of oxygen-sensitive transcription factors.

## Acknowledgments

We thank H. Zhao, D. Lambrechts, and P. Carmeliet (Vesalius Research Center, Belgium) for sharing their TCGA tumor annotations. We thank R. Looper (U. of Utah) for synthesizing and sharing the esterified 2-HG. M. K. Koski and R. Wierenga (U of Oulu, Finland) for help with structural modelling, and T. Aatsinki and E. Lehtimaki (U of Oulu, Finland) for technical assistance. We thank the Broad GDAC group for help with the GDAC-GSEA source code. We thank M. Oser for sharing mouse KDM5A sgRNAs and V. Koduri for assistance in acquiring microscopic images.

## Author Contributions

A.A.C. and W.G.K. conceived experiments and wrote the manuscript, A.A.C, T.L., M.M., A.E.R., A.C., R.J., and J.S. performed experiments. A.A.C., T.L., M.M, A.E.R, M.A.B, M.Y.T, Y.J.M., S.M., S.K.M, B.O., Z.T.H, S.M., J.J., S.S., M.H., R.B., P.K., and W.G.K. analyzed data.

## Data and materials availability

All data is available in the main text or the supplementary materials. RNA-Seq and ChIP-Seq data was uploaded to the GEO (GSE114086).

## List of Supplementary Materials

Materials and Methods

Figures S1-S25

Tables S1-S6

References (61-76)

