## Supplemental Methods and Data for "The UTX Tumor Suppressor Directly Senses Oxygen to Control Chromatin and Cell Fate"

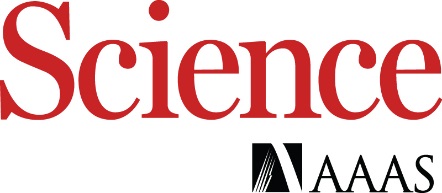

Supplementary Materials for

**Title: The UTX Tumor Suppressor Directly Senses Oxygen to Control Chromatin and Cell Fate**

**Authors:** Abhishek A. Chakraborty, Tuomas Laukka, Matti Myllykoski, Alison E. Ringel, Matthew A. Booker, Michael Y. Tolstorukov, Yuzhong Jeff Meng, Sam Meier, Rebecca B. Jennings, Amanda L. Creech, Zachary T. Herbert, Jessica Spinelli, Samuel K. McBrayer, Benjamin A. Olenchock, Jacob D. Jaffe, Marcia C. Haigis, Rameen Beroukhim, Sabina Signoretti, Peppi Koivunen, and William G. Kaelin, Jr.

Correspondence to:

W.G.K or P.K.

**This PDF file includes:**

Materials and Methods

Figs. S1 to S25

**Other Supplementary Materials for this manuscript include the following:**

Tables S1 to S6

MATERIALS AND METHODS

**Cell lines**

The C2C12 cells (CRL-1772), MCF7 cells (HTB-22), and 293T cells (CRL-3216) were obtained from ATCC. The mHepa-1 c4 cells (a gift of Dr. Oliver Hankinson) were maintained in Minimum Essential Medium (MEM) alpha with Earle’s salts supplemented with 10% fetal bovine serum (FBS) (Life Technologies 10437-028) and 1X penicillin-streptomycin (Life Technologies 15140163). C2C12 and RCC4 cells (a gift of Dr. Peter Ratcliffe) were maintained in Dulbecco’s Modified Eagle’s Medium (DMEM) (Life Technologies) supplemented with 10% FBS and 1X penicillin-streptomycin. The 293T cells were maintained in Dulbecco’s Modified Eagle’s Medium (DMEM) (Merck) supplemented with 10% FBS, 2 mM L-glutamine (Sigma G7513), 0.375 % Na-bicarbonate (Biowest L0680), and 1X penicillin-streptomycin (Sigma P0781). The 293T media supplemented with 10 µg/ml insulin was used to culture MCF7 cells. The SK-N-BE(2) cells, a gift ftom Dr. Lucy Godley at University of Chicago, were maintained in RPMI1640 media (ThermoFisher). The glioma cell lines TS603 and TS516 (a gift of Dr. Ingo Mellinghoff at MSKCC) (*61*) and the BT260 cells (a gift of Dr. Keith Ligon at DFCI) (*62*), were cultured in NeuroCult NS-A Basal Medium with Proliferation Supplement (StemCell Technologies) supplemented with EGF (20 ng/ml), bFGF (20 ng/ml), heparin (2 μg/ml), 1% penicillin/streptomycin, Fungizone (250 ng/ml), and Plasmocin (2.5 μg/ml). The glioma cell line MGG152 (a gift of Dr. Daniel Cahill at MGH) (*63*), was cultured in Neurobasal Medium supplemented with 3 mM glutamine, 1X B27, 0.25X N2, EGF (20 ng/ml), bFGF (20 ng/ml), heparin (2 μg/ml), 1% penicillin/streptomycin, and Fungizone (250 ng/ml). All glioma cell lines were passaged as neurospheres and dissociated 1-2 times per week. The Mouse Embryonic Fibroblasts expressing MyoD-ER, a kind gift from Dr. Benevolenskaya (U. of Illinois, Chicago) (*64*), were cultured in DMEM supplemented with 10% Calf Serum and 1X Penicillin/Streptomycin. The primary Human Skeletal Myoblasts (HSkM) were purchased from Gibco (A12555) and cultured in Complete Skeletal Muscle Media (ZenBio). The human mammary epithetelial cells (HMLE) were a kind gift from Dr. William Hahn and were cultured in MEBM basal Medium (Lonza CC-3151) supplemented with growth factors (MEGM Singlequot, Lonza CC-4136). Lentivirally infected cells were selected with puromycin (2 µg/ml) or blasticidin (10 µg/ml) as appropriate for the vector used. All mammalian cells were grown at 37°C in 5% CO_2_. The *Sf9* insect cells were cultured in TNM-FH media supplemented with 10% FBS and cultured at 27°C.

**Plasmids**

The pLX304-GFP (ccsbBroad304_99986) vector was obtained from the Gene Pertubation Platform (GPP, Broad Institute). The pLX304 ARNT expression plasmids were made by sub-cloning the human ARNT cDNA (wild-type or with a 5’ 414 bp deletion [Δ414]) generated by PCR amplification into the pLX304 destination vector (Addgene # 25890) using Gateway cloning. The pLKO-based plasmids to express shRNAs targeting *Kdm6a* (, #1: TRCN0000107762 and #2: TRCN0000107763) and *Kdm6b* (, #1: TRCN0000236677 and #2: TRCN0000236678) were obtained from the GPP, Broad Institute.

The baculovirus for C-terminal FLAG-tagged human KDM5B was a gift from Dr. Qin Yan at Yale University. Human full length KDM4A and KDM4B in the pReceiver-I01 plasmid, a gift from Dr. Susanne Schlisio at Karolinska Institute, were used to generate the corresponding baculoviruses with N-terminal His-tags. FLAG-tagged human full length KDM5A, KDM6A, KDM6B, and the JumonjiC-domains of KDM6A (931→End) and KDM6B (1164→End) were generated by PCR and subcloned into the pVL1393 backbone. The KDM6A M→T/E→D mutant was generated via site-directed mutagenesis of pVL1393-KDM6A-FLAG using the QuickChange® Lightning site-directed mutagenesis kit (Stratagene) according to the manufacturer's protocol and confirmed by Sanger DNA sequencing. The corresponding baculoviruses with C-terminal FLAG-tags were generated using the BacMagic-3 DNA kit (Novagen). The FLAG-tagged full-length human KDM5C and KDM5D were generated by PCR and subcloned into pFastBac Dual plasmid. KDM5C and KDM5D bacmids were generated using DH10Bac cells with the standard Bac-to-Bac protocol (Invitrogen) and the corresponding baculoviruses were generated by transfecting the bacmid DNA into Sf9 cells.

The pLX304-WT KDM6A expression vector was made by subcloning a human KDM6A cDNA generated by PCR amplification of pCMV-HA-UTX (Addgene#24168) into the pLX304 destination vector by Gateway cloning. pLX304 expression plasmids encoding UTX/KDM6A mutants were made by site-directed mutagenesis (Quickchange II XL; Agilent 200521) of pLX304-KDM6A according to the manufacturer’s recommendations and confirmed by Sanger DNA sequencing. The pLenti-CRISPRv2 (Addgene # 52961) sgRNA expression plasmids were made by cloning annealed duplex oligonucleotides (see table S1) into BsmBI-digested pLenti-CRISPRv2. All primer sequences used for cloning, mutagenesis, and Real-Time qPCR analysis are provided in the table below.

**Histone extraction for immunoblotting and mass spectrometry profiling**

To prepare crude histone extracts for immunoblotting, soluble subcellular fractions were eliminated by lysing cells twice in nucleus buffer (15 mM Tris [pH 7.5], 250 mM Sucrose, 60 mM KCl, 15 mM NaCl, 5 mM MgCl_2_, and 1 mM CaCl_2_) supplemented with 0.5% NP-40, 1 mM DTT, 10 mM sodium butyrate, and protease inhibitor cocktail (Roche, 29384100). Soluble fractions were discarded by centrifugation at 8000 rcf for 5 min at 4°C. The insoluble (chromatin-enriched) fractions were acid extracted with 0.2N HCl for >6 hours (typically overnight) at 4°C. For immunoblot analysis the histone preparations were quantified using the Protein Assay Dye Reagent (Biorad 500-0006) and 200 ng histone from each sample was analyzed per lane.

For the Global Chromatin Profiling (GCP) assay histone preparations were further purified by TCA precipitation. The precipitated histones were derivatized with NHS-propionate, digested by trypsinization, rederivatized, and modification profiles were generated by mass spectrometric comparison to a spiked-in internal standard peptide for the modification of interest as previously described (*35*).

**Immunoblot and Immunoprecipitation analysis**

Cells were lysed in lysis buffer (50 mM Tris HCl [pH 7.5], 400 mM NaCl, 1% Nonidet P-40, 1 mM EDTA, and 10% glycerol) supplemented with a protease inhibitor cocktail (PIC, Complete mini EDTA-Free, Roche) and a phosphatase inhibitor cocktail (PhosSTOP, Roche). For immunoprecipitation analysis, 750 µg of total protein was diluted to a total volume of 1 ml, of which 10% (100 µl) was kept aside as “Input”, and the remainder was tumbled in the presence of 30 µl 50% anti-V5 sepharose (Sigma A7345) for 3 hours at 4°C. Bound beads were washed 4 times in lysis buffer, boiled in 100 µl of 1X SDS buffer (0.0005% Bromophenol Blue in 50 mM Tris HCl [pH 6.8], 0.15% β-Mercaptoethanol, 2% SDS, and 10% Glycerol), and resolved by SDS-PAGE. For immunoblot analysis, lysates prepared in 1X SDS buffer were resolved using 6%, 8%, or 15% SDS-PAGE gels, as appropriate, and transferred onto 0.2 µM nitrocellulose membranes.

The following primary antibodies were used for immunoblot analysis: Actin (Cell Signaling 4970, 1:5000), ARNT (Cell Signaling 5537, 1:1000), Cyclin E (Cell Signaling 4129, 1:1000), EZH1 (Novus 56358), EZH2 (Cell Signaling 5246, 1:1000), HIF1α (Cell Signaling 14179, 1:1000), Histone H3K27me3 (Cell Signaling 9733, 1:2000), Histone H3K4me3 (Cell Signaling 9727, 1:2000), Histone H3K9me3 (Abcam ab8898, 1:2000), Histone H3 (Cell Signaling 4499, 1:2000), KDM5A (Abcam 70892, 1:1000), KDM6A (Cell Signaling 33510, 1:1000), KDM6B (Abcam 38113, 1:1000), MyHC (Sigma MY32, 1:5000), MYC (Cell Signaling 5605, 1:1000), Tubulin (Sigma T5168, 1:5000), and Vinculin (Sigma V9131, 1:5000). HRP-conjugated secondary antibodies (Pierce, 1:5000) targeting the primary antibodies were detected with and chemiluminescent HRP substrates (Supersignal West Pico, Thermo Fisher Scientific; Supersignal West Femto, Thermo Fisher Scientific; or Immobilon, Millipore).

**Metabolite Profiling**

To measure changes in intracellular metabolite levels, cells were seeded in 6-well plates in their respective growth or differentiation media, as appropriate. To isolate metabolites the cells were placed on ice and the cell culture was removed by aspiration. The cells were then washed twice with ice cold saline and snap frozen by floating the cell culture plates on liquid Nitrogen. To extract metabolites, frozen cells from each well were scraped in 400 µl of 75% Methanol and transferred into fresh Eppendorf tubes. 300 µl of Chloroform was then added and vortexed at 4°C for 20 min. Samples were centrifuged at 16.2K rcf for 10 min at 4°C on an Eppendorf 5415R tabletop centrifuge and 200 µl of the aqueous (top) layer was collected and dried overnight at 4°C in a CentriVap vacuum concentrator (Labonco). Dried samples were derivatized by first resuspending in 20 µl Methoxamine (MOX) and incubating for 60 min at 37°C, followed by the addition of 30 µl N-tert-butyldimethylsilyl-N-methyltrifluoroacetamide (TBDMS) and incubation for 90 min at 65°C. Derivatized metabolites were analyzed on an Agilent 7890B GC/5977A MSD system.

Enantiomer-specific 2-HG measurements were performed by LC-MS using a modified version of a previously published method (*30*). Cell lysates were extracted in a mixture of 75% methanol and chloroform spiked with 0.1 µg/ml of racemic mixture of ^13^C_5_-2HG as an internal standard, and dried overnight at 4°C in a CentriVap vacuum concentrator (as described in detail above). Metabolites were resuspended in 100 μl of 50 mg/ml diacetyl-L-tartaric anhydride (DATAN, Sigma 358924) prepared fresh in a solution of dichloromethane (Sigma 650463) and acetic acid (Sigma 45754) [v/v = 4:1] and derivatized at 70°C for 2 hours. Derivatized samples were dried in a speedvac for 1 hour, resuspended in 50 μl of UltraPure water (18.2 MΩ, PureLab), and clarified by centrifugation at 4°C. 5 µl was analyzed by LC–MS/MS.

Chromatographic separation was performed using an Agilent Infinity 1290 II LC system, and detection was performed using an Agilent 6470 triple-quadrupole mass spectrometer operating in MRM and negative ion modes with Jet Stream source. Samples were ionized with the following MS source parameters: nebulizer = 45 psi, capillary voltage = 2000 V, nozzle voltage = 500 V, sheath gas temperature = 325°C, sheath gas flow = 12 L/min., gas flow = 13 L/min., and gas temperature = 150°C. The fragmentor energy was set at 70 V and the cell accelerator voltage was set at 4 V for all compounds. MRM transitions were monitored for DATAN-derivatized compounds using the following parent ion, fragment ions, and collision energies (CE), with the primary transition for ion quantification indicated using an asterisk: ^13^C_5_-2HG m/z 368 🡪 134* (CE = 29 V) and m/z 360 🡪 152 (CE = 5 V), 2HG m/z 363 🡪 129* (CE = 29 V) and m/z 363 🡪 147 (CE = 5 V), malate m/z 349 🡪 133* (CE = 14 V) and m/z 349 🡪 115 (CE = 28 V), and lactate m/z 305 🡪 89.1 (CE = 14 V).

Enantiomer identities were assigned using the retention time for the ^13^C-labeled internal standard added to each sample. Chromatograms were analyzed using Agilent MassHunter Quantitative Analysis (for QQQ) software.

For quantification of absolute levels of intracellular 2-HG by GC-MS, total ion counts measured from the respective samples was quantified against a standard curve of L-2HG in water. Absolute levels of enantiomer-specific 2-HG were calculated either by normalizing the peak area measured by LC-MS either against an external standard curve prepared in the same cellular matrix or using the peak areas recorded for the ^13^C_5_-2HG internal standard. Pilot experiments confirmed no measurable difference in quantifications obtained using either approach. For both GC-MS and LC-MS based approaches, we assumed that all suspended cells were perfect spheres. Cell counts and average cell diameter in each well were used to define the total intracellular volume {Volume = [ πx(Average Cellular Diameter)^3^]/6}, thus allowing for an absolute quantification of intracellular 2-HG concentration per well.

**Baculoviral Protein Expression and Purification**

Recombinant proteins were produced by transducing *Sf9* insect cells with the corresponding baculoviruses for 72 h. The cells were then washed with cold 1X PBS, and homogenized in a buffer containing 10 mM Tris-HCl pH 7.8, 150 mM NaCl, 100 mM glycine, 5 *µ*M FeSO_4_, 0.1% Triton X-100 and PIC. The soluble fractions of the FLAG-tagged enzymes were affinity purified using the anti-FLAG M2 affinity gel (Sigma), washed with TBS containing 5 *µ*M FeSO_4 and_ PIC, and eluted with 150 µg/ml FLAG-peptide. KDM6B (1164→End) was further purified using UNOQ1 anion exchange column (Bio-Rad) to eliminate nucleic acid contamination. KDM6B was bound to the column in the FLAG affinity gel elution buffer and eluted with a gradient from 0 to 1 M NaCl in 50 mM Tris-HCl [pH 7.6]. For His-tagged KDM4A and KDM4B, the soluble fractions were affinity purified using Ni-NTA agarose (Qiagen) and eluted with imidazole, which was then removed with PD-10 columns (Sigma).The fractions collected were analyzed using 10% SDS-PAGE under reducing conditions followed by Coomassie Blue staining.

**Circular dichroism (CD) spectroscopy**

CD spectroscopy was performed using a Chirascan CD spectrometer (Applied Photophysics, Leatherhead, UK). CD data were collected between 190 and 260 nm at 22°C using a 1 cm path-length quartz cuvette. CD measurements were acquired every 1 nm with 1 s as an integration time and repeated three times with baseline correction. The data were analyzed with Pro-Data Viewer (Applied Photophysics).

**Determination of melting temperatures**

Thermal unfolding of wild-type and MT/ET KDM6A were recorded between 210 and 260 nm from 22°C to 94°C with a 2°C step size at 1°C/min ramp rate with ± 0.2°C tolerance. The melting temperatures were analyzed with Global3 (Applied Photophysics).

**Enzyme kinetic assays**

Oxygen *K_M_* values for the KDMs were measured using a modified version of a previously described assays (*8*) (*65*), based on the stoichiometric coupling of lysine demethylation to 2-OG decarboxylation. Each 50 µl reaction volume consisted of 50 mM Tris-HCl [pH 7.8] (except for KDM5A, which was in MES [pH 6.5], and KDM5C and KDM5D, which were in Tris-HCl [pH 7.5]), 2 mg/ml bovine serum albumin (Roche), 60 µg/ml catalase (Sigma), 0.1 mM dithiothreitol, 2 mM sodium ascorbate, 50 µM iron(II) sulfate, 10% (v/v) dimethyl sulfoxide, and 50 µM (KDM4A and KDM4B), 40 µM (KDM5A, KDM5C and KDM5D), or 200 µM (KDM5B, KDM6A and KDM6B) of 2-oxo[1-^14^C]glutarate (Perkin-Elmer). Respective histone peptides were used as substrates at saturating concentrations. These were 200 µM and 500 µM histone H3(1-19)K9me3 (Innovagen) for KDM4A and KDM4B, respectively, 30 µM (KDM5A), 20 µM (KDM5B), 15 µM (KDM5C and KDM5D) histone H3(1-21)K4me3 (Anaspec or Innovagen), respectively , and 1000 µM and 100 µM histone H3(21-44)K27(me3) (Innovagen) for KDM6A and KDM6B, respectively. All peptide substrates contained an additional glycine and a biotinylated lysine residue at their C-termini. Enzyme concentrations were 0.2-1.5 μM. Enzymatic reactions were simultaneously performed in five to six different oxygen concentration using an InVivo400 hypoxia workstation (Ruskinn), following an initial reagent equilibration at the corresponding oxygen concentration. The reaction endpoints, chosen within the linear phase of the enzymatic assay time curves (*38*) and Fig. S7), were the following: KDM4A, KDM4B, KDM5B, KDM6A and KDM6A (931→End) 20 min, KDM6B (1164→End) 15 min, Fig KDM5A, KDM5C, KDM5D and KDM6B 3 min. Reactions were stopped by adding 100 µl of 1M potassium phosphate pH 5 and the amount of ^14^C-labeled CO_2_ generated was scintillated in a Tri-Carb 2900TR (Perkin Elmer). When the effect of H_2_O_2_ on the catalytic activity of KDM6A and the MT/ED mutant was studied catalase was omitted from the reaction. *K_M_* values were determined from the Michaelis-Menten saturation curves and Lineweaver-Burk plots using Excel (Microsoft). *V_max_* values for wild type KDM6A and the KDM6A MT/ED mutant were calculated from the Michaelis-Menten saturation curves and Lineweaver-Burk plots using Excel (Microsoft) after standardizing observed d.p.m. values for enzyme amount quantified from the Coomassie Blue-stained SDS-PAGE gel. L-2HG IC_50_ values for wild-type and MT/ED KDM6A were determined by increasing the concentration of L-2HG in the presence of non-limiting amounts of other factors, and in the presence of 37.5 μM 2-oxoglutarate.

**Myogenic Differentiation Assays**

C2C12 cells were seeded in 6-well plates in growth medium [DMEM, 10% FBS, and 1X Penicillin/Streptomycin] at a density of 200,000 cells/well and allowed to adhere overnight at 37°C. Differentiation was induced for 4 days by switching cells into differentiation media [DM: DMEM, 2% Horse serum, 1X insulin-transferrin-selenium (ITS)] supplemented with 1X Penicillin/Streptomycin on day 0, followed by media changes on day 2 and day 3. For Metformin rescue experiments, cells were seeded in growth medium in the presence of the indicated concentrations of Metformin, and Metformin treatment was continued in differentiation medium throughout the course of the experiment.

Mouse Embryonic Fibroblasts lentivirally transduced to express a MyoD-ER fusion were cultured and seeded (as described above) in growth medium [DMEM, 10% FBS] supplemented with 1X Penicillin/Streptomycin and then induced to differentiate by being switched to DM using the schema described in fig. S12.

Differentiation assays were scored by measuring the mRNA levels of myogenic differentiation markers by Real-Time qPCR and by scoring “Fusion Index (FI)”. FI was measured by counting multi-nucleated MyHC positive myotubes from 3 representative fields (~350 cells/field) from each biological replicate. Mean FI (±SD) from 3 biological replicates was compared by statistical analysis.

Human Skeletal Myoblasts were seeded at a density of 300,000 cells per well of a 6-well plate in Skeletal Muscle Growth Medium (ZenBio, SKM-M) and allowed to adhere overnight at 37°C. Differentiation was induced in DM for 3-4 days with daily media changes.

**Immunofluorescence staining**

For immunofluorescence assays, C2C12 cells were seeded on coverslips and cultured in 6-well plates under the desired conditions. To fix cells, the media was aspirated and cells were washed twice with cold 1X PBS, and fixed using 2% formaldehyde in 1X PBS for 15 minutes at room temperature. Cells were permeabilized on ice for 5 mins using 0.5% TritonX-100 in 1X PBS and then blocked for 10 minutes using blocking buffer [5% normal goat serum (NGS) in 1X PBS]. Cells were stained sequentially with primary and secondary antibody diluted in wash buffer [1% normal goat serum (NGS) in 1X PBS] for 1 hour each at room temperature. Cells were washed thrice after both primary and secondary antibody staining. The cells were then counterstained with DAPI and mounted onto slides using mounting medium. Stained cells were imaged using a Nikon inverted TE2000 inverted fluorescence microscope equipped with a motorized platform and a Hamamatsu Orca ER digital CCD camera and analyzed using ImageJ.

**Immunohistochemistry**

For immunohistochemistry analyses, 4 um thick paraffin-embedded tissue sections were prepared and left to air-dry overnight. Slides were baked in an Isotemp Oven (Fisher Scientific) for 30 minutes at 60°C to melt excess paraffin. Immunohistochemical staining for H3K27me3 was performed on a Bond III automated stainer (Leica Biosystems) using the Bond Polymer Refine Detection Kit (Leica Biosystems cat. no. DS9800). Briefly, antigen retrieval was performed using the Bond Epitope Retrieval Solution 1 (Citrate, pH 6.0) for 30 minutes, following which slides were incubated with a rabbit monoclonal antibody directed against H3K27me3 (Cell Signaling Technology cat. no. 9733, 1:200) diluted in Bond Primary Antibody Diluent (Leica Biosystems cat. no. AR9352) for 30 minutes. Slides were subsequently incubated with the HRP-conjugated secondary antibody for 10 minutes. Staining was visualized by incubating the slides with the chromogen 3,3’-diaminobenzidine for 5 minutes. Finally, slides were counterstained with hematoxylin, dehydrated in graded ethanol and xylene, and cover-slipped. Typically 3 serial sections from 2 male and 2 female mice (for main figure 2) and 5 independent tumors (for fig. S10) were stained and analyzed.

**Flow cytometry analysis**

Flow cytometry analysis for intracellular Myosin Heavy Chain expression was performed on cells fixed for 15 minutes with 1% Formaldehyde at room temperature. Cells were permeabilized on ice for 10 mins using 0.5% TritonX-100 in 1X PBS and then blocked for 30 minutes in blocking buffer. Cells were stained with primary antibody (APC-conjugated anti-MyHC, R&D Biosystems) for 3 hours at room temperature in the dark. For intracellular ROS measurements, cells were cultured for 2 hours in the presence of deep red CellROX (Invitrogen), as per the manufacturer’s instructions. The cells were then harvested using trypsin and resuspended in growth medium. For CD44 staining, HMLE cells fixed as above were washed once in ice-cold PBS and resuspended in 50 µl of staining solution [5 µl AlexaFluor 700 conjugated anti-CD44 (BD 561289), 1% FBS, in 1X PBS] and incubated at 25°C in the dark for 2 hours. For all measurements, cells were washed thrice, resuspended in wash buffer (1%FBS in 1X PBS), and then analyzed by flow cytometry using a BD LSR II system. In all flow based assays, measurements from 10,000 cells were collected for analysis and measurements from 1000 cells was displayed in the plots.

**RNA-Seq and transcriptional analysis**

RNA-Seq analysis was performed on total RNA extracted from cells using Trizol (Life Technologies) according to the manufacturer’s instructions. Total RNA was sent to the Molecular Biology Core Facility (MBCF, Dana-Farber Cancer Institute) for library construction and sequencing. Libraries were prepared from 500ng of purified total RNA using Illumina Truseq Stranded mRNA samples preparation kits according to the manufacturer’s protocol. Finished libraries were quantified by Qubit fluorometer, Agilent TapeStation 2200, and RT-qPCR using the Kapa Biosystems library quantification kit according to manufacturer’s protocols. Uniquely indexed libraries were pooled in equimolar ratios and sequenced on an Illumina NextSeq500 run with single-end 75bp reads.

Sequenced reads were aligned to the UCSC mm9 reference genome assembly and gene counts were quantified using STAR (v2.5.1b) (*66*) and normalized read counts (RPKM) were calculated using cufflinks (v2.2.1) (*67*). RNAseq analysis was performed using the VIPER snakemake pipeline (*68*).

**ChIP-Seq Analysis**

ChIP-Seq analysis was performed using C2C12 cells cultured under the desired conditions. Briefly, after aspirating culture medium, cells were washed with cold 1X PBS, and fixed using 1% formaldehyde (reconstituted in 1X PBS) at room temperature for 10 min. Excess formaldehyde was quenched by dropwise addition of Glycine to a final concentration 125 mM and further incubation at room temperature for 5 min. Fixed cells were harvested, washed 1X with cold 1X PBS, lysed in SDS buffer (50mM Tris-HCl [pH 8], 1% SDS, 10mM EDTA), and sonicated in the cold using a microprobe (5 cycles: 10” on/10” off). Lysates were clarified by centrifugation at 16.2K rcf for 10 min at 4°C. To test for sonication efficiency and quantify the concentration of chromatin, 20 µl of the clarified lysates was aliquoted into a fresh tube. Sample volume was adjusted to 100 µl using TE, and treated sequentially, first by addition of 1 µl RNAse (10 mg/ml, Roche) for 30 min at 37°C, followed by addition of 5 µl of Proteinase K (20 mg/ml) and incubation overnight at 65°C. DNA was recovered by column purification using a Qiagen kit, as per the manufacturer’s instructions, run on a gel to ensure an average size of <500bp, and quantified on a Nanodrop.

ChIP was performed in a 1000-1500 µl reactions reconstituted in ChIP Dilution Buffer [20mM Tris-HCl pH 8, 1% Triton-X 100, 2mM EDTA, 150mM NaCl + PIC], diluting SDS Buffer at least 1:5 (ideally 1:10), and using 3 µg chromatin, 10 µl of anti-H3K27me3 (Cell Signaling), and 50 µl of blocked A/G beads.

Blocked A/G beads were prepared in advance by incubating 500 µl of 50% slurry (1:1 Protein A:Protein G beads) with 15 µl Sonicated Herring Sperm DNA (10 mg/ml) and 50 µl BSA (10 mg/ml), overnight at 4°C. Beads were washed thrice with 1 ml cold ChIP Dilution Buffer and re-adjusted to 50% slurry by addition of 250 µl of ChIP Dilution Buffer.

ChIPs were performed by tumbling down the reconstituted reactions for 3 hours are 4°C. Beads were washed thrice with ChIP Dilution Buffer and bound DNA was recovered from the beads by following the RNAse and ProteinaseK treatments, followed by purification over the Qiagen column, as described above. Recovered DNA was quantified by Qubit and used to prepare libraries using the Illumina Nextera library prep method. Samples were run on a NextSeq 500 and mapped and analysed using the ChiLin analytical pipeline (*69*). Genomic H3K27me3 recruitment was compared using the Integrated Genome Viewer (*70*).

ChIP-Seq reads generated from H3K4me3 and H3K27me3 libraries were aligned to the mouse genome (mm9) using bowtie2 (*71*) using the default options. Enrichment of the histone marks were calculated relative to input in 50-bp non-overlapping bins using the R package spp (*72*). Tag density was estimated using a Gaussian kernel with bandwidth of 35 as described in spp package. Standard BigWig files were generated for both tag enrichment and tag density for visualization in the genome browser. Reference transcript annotations for the mouse genome were obtained from the University of California Santa Cruz Genomics Institute (UCSC) (*73*). Mean histone enrichment was calculated within a -2Kb to +5Kb region relative to each transcription start site (TSS). For gene bodies, mean histone enrichment was calculated from the start site of each transcript to the termination site. Mean histone enrichment was then compared to expression data produced by the VIPER package (*68*) and correlations were calculated using R. Gene ontology enrichment for H3K4me3 or H3K27me3 marked genes (normoxia versus hypoxia) was carried out from cells in differentiation media using PANTHER (*74, 75*), employing a TSS-proximal enrichment fold-change cutoff of >=2 and a significance cutoff of p < 0.05.

**Gene Set Enrichment Analysis**

Gene set enrichment analysis (GSEA) (*45*) was performed using TCGA data on gene sets in the Hallmark collection (MSigDB <http://software.broadinstitute.org/gsea/msigdb>) and curated gene sets related to H2K27me3 (Table. S2). Tumors that were previously annotated as “hypoxic” or “normoxic” based on HIF signature (*6*) (Table. S1), were analyzed from the TCGA cohorts BLCA, BRCA, COAD, HNSC, LUAD, LUSC, and UCEC. Standardized RSEM RNA-seq data was downloaded from the Broad Institute TCGA GDAC Firehose repository (*76*). Samples sequenced by Illumina Genome Analyzer and HiSeq machines were pooled and duplicate samples were removed.

GSEA parameters are as follows:

  --GSEA.v.1.0.reshuffling.type "sample.labels" \

  --GSEA.v.1.0.nperm 500 \

  --GSEA.v.1.0.weighted.score.type 1 \

  --GSEA.v.1.0.nom.p.val.threshold -1.0 \

  --GSEA.v.1.0.fwer.p.val.threshold -1.0 \

  --GSEA.v.1.0.fdr.q.val.threshold 0.25 \

  --GSEA.v.1.0.topgs 20 \

  --GSEA.v.1.0.adjust.FDR.q.val F \

  --GSEA.v.1.0.gs.size.threshold.min 25 \

  --GSEA.v.1.0.gs.size.threshold.max 500 \

  --GSEA.v.1.0.reverse.sign F \

  --GSEA.v.1.0.perm.type 0 \

**Structural analyses**

The crystal structure coordinates for KDM6A (PDB: 3avr) and KDM6B (PDB: 2xue) demethylase domains (*53, 54*) were downloaded from Protein Data Bank. The structures were aligned and a composite image of the active site was created in PyMOL. Atomic distances between residues lining the catalytic pocket were measured using PyMOL.

**Respiration Measurements**

Respiration was assessed using the Seahorse XFe-96 Analyzer (Seahorse Bioscience). C2C12 cells were cultured in differentiation medium with the desired concentration of metformin in either 2% or 21% oxygen for 2 days prior to respiratory measurements. On the day of the measurement, differentiation medium was aspirated and cells were pretreated for 1 hour with serum-free Seahorse Media (Seahorse Bioscience, 102353) supplemented with 10 mM glucose, 2 mM L-glutamine, and 1 mM sodium pyruvate, at their respective oxygen conditions and at the appropriate metformin concentration. Basal oxygen consumption rate (OCR) was measured in 3 minute intervals over a period of 30 minutes. To normalize for cell density, basal OCR was normalized to protein content. Cells were lysed in each well with RIPA buffer (10 mM Tris-Cl [pH 8], 140 mM NaCl, 1 mM EDTA, 1% Triton X-100, 0.1% sodium deoxycholate, 0.1% SDS, and 1 mM PMSF) and protein concentrations were measured using a BCA assay (Thermo Scientific, 23227).

**Statistical Analysis**

All experimental results represent observations from at least 3 biological replicates, except for the genomics data (ChIP-Seq and RNA-Seq), which were measured from 2 biological replicates. All genomic analysis were performed in R using the “spp” package (*72*), as described in the methods. Unless indicated otherwise, all other data are presented as Mean ± SD and the number of replicates is indicated in the legends. All statistical analysis were performed using the GraphPad Prism package. Statistical significance was calculated by *t-test*, assuming gaussian distribution, unpaired with the Welch’s correction, and values below 0.05 were considered significant. Multiple testing corrections, if necessary, were performed by the Holm-Sidak method using Graphpad Prism. For GSEA (*45*), as described in the algorithm’s manual, a FDR (q) value below 0.25 was considered significant.

**Primer/Oligos List**

| **No.** | **Name** | **Sequence** | **Remarks** |
| --- | --- | --- | --- |
| 1 | ARNT (*68*)F | ggggacaactttgtacaaaaaagttggcATGGCAGCGACTACTGCCAA | Primers used for gateway sub-cloning wild-type of mutant (414) ARNT into the pLX304 vectors |
| 2 | ARNTΔ414(*68*)F | ggggacaactttgtacaaaaaagttggcATGAAGTCCTTGCGGGGAAC |  |
| 3 | ARNT (*68*)R | ggggacaactttgtacaagaaagttgggcaTTCTGAAAAGGGGGGAAACAT |  |
| 4 | mNdrg1_F | ATGTCCCGAGAGCTACATGA | Primers used for Real-Time qPCR analysis |
| 5 | mNdrg1_R | TGACATGCAGGGAGCCATGT |  |
| 6 | mAdm_F | CCTTCGCAGTTCCGAAAGAA |  |
| 7 | mAdm_R | AGTTGTGTTCTGCTCGTCCA |  |
| 8 | mEgln3_F | CTATGTCAAGGAGCGGTCCAA |  |
| 9 | mEgln3_R | GTCCACATGGCGAACATAACC |  |
| 10 | mKDM6A_F | TACGAATCTCTAATCTTAAA |  |
| 11 | mKDM6A_R | TTCCAGTAATCAGACTGTAA |  |
| 12 | mKDM6B_F | CCTGCAGTCAATGAAGCACTG |  |
| 13 | mKDM6B_R | CTCCACGTCGCATTCGTTG |  |
| 14 | ExoUTX_F | TTGATCTGCTTTTTGTCACT |  |
| 15 | ExoUTX_R | ACCGAGGAGAGGGTTAGGGAT |  |
| 16 | mActin_F | TAGGCACCAGGGTGTGATG |  |
| 17 | mActin_R | CATGGCTGGGGTGTTGAAGG |  |
| 18 | 6Asg1Top | CACCGTCCTTGGCTCGACAAAAGCT | Oligos used to generate sgRNAs targeting *Kdm6a* |
| 19 | 6Asg1Btm | AAACAGCTTTTGTCGAGCCAAGGAC |  |
| 20 | 6Asg2Top | CACCGCCGCCTTTTCGGGTTCGTG |  |
| 21 | 6Asg2Btm | AAACCACGAACCCGAAAAGGCGGC |  |
| 22 | ARNTsg1Top | CACCGGGCTATTAAGCGACGGTCA | Oligos used to generate sgRNAs targeting *Arnt* |
| 23 | ARNTsg1Btm | AAACTGACCGTCGCTTAATAGCCC |  |
| 24 | ARNTsg2Top | CACCGAGAAACGGCCATGCGTAAGA |  |
| 25 | ARNTsg2Btm | AAACTCTTACGCATGGCCGTTTCTC |  |
| 26 | EZH2sg1Top | CACCGCGGCCCCCTGGGCGTTTAGG | Oligos used to generate sgRNAs targeting *EZH2* |
| 27 | EZH2sg1Btm | AAACCCTAAACGCCCAGGGGGCCGC |  |
| 28 | EZH2sg2Top | CACCGAATAACTGCACTTACGATGT |  |
| 29 | EZH2sg2Btm | AAACACATCGTAAGTGCAGTTATTC |  |
| 30 | 6A_MT_Top | Aatttgaatttcctaacgggttcttggtggcccaa | Oligos used for site-directed mutagenesis of KDM6A cDNA |
| 31 | 6A_MT_Btm | ttgggccaccaagaacccgttaggaaattcaaatt |  |
| 32 | 6A_ED_Top | tactgtagcatttgtgatgtggaggtttttgat |  |
| 33 | 6A_ED_Btm | atcaaaaacctccacatcacaaatgctacagta |  |
| 34 | 6Bsg1Top | CACCGAAGCTTCCTCCATAGCGAA | Oligos used to generate sgRNAs targeting *Kdm6b* |
| 35 | 6Bsg1Btm | AAACTTCGCTATGGAGGAAGCTTC |  |
| 36 | 6Bsg2Top | CACCGCTGCAAGCGGCCAATCCG |  |
| 37 | 6Bsg2Btm | AAACCGGATTGGCCGCTTGCAGC |  |
| 38 | Hu6Asg1Top | CACCGCAGCATTATCTGCATACCAG | Oligos used to generate sgRNAs targeting *KDM6A* |
| 39 | Hu6Asg1Btm | AAACCTGGTATGCAGATAATGCTGC |  |
| 40 | Hu6Asg2Top | CACCGTTGGATAATCTTCCAATAAG |  |
| 41 | Hu6Asg2Btm | AAACCTTATTGGAAGATTATCCAAC |  |
| 42 | mKdm5A sg1top | CACCGTCTTTGAGCCCAGTTGGG | Oligos used to generate sgRNAs targeting *Kdm5a* |
| 43 | mKdm5A sg1btm | AAACCCCAACTGGGCTCAAAGAC |  |
| 44 | mKdm5A sg2top | CACCGGCGCCCGATAAAACTCAG |  |
| 45 | mKdm5A sg2btm | AAACCTGAGTTTTATCGGGCGCC |  |
| 46 | L2HGDHsg1top | CACCGAAAGAAGGAGCCGTATTGCA | Oligos used to generate sgRNAs targeting *L2hgdh* |
| 47 | L2HGDHsg1btm | AAACTGCAATACGGCTCCTTCTTTC |  |
| 48 | L2HGDHsg2top | CACCGACCTCAAGGGAATTCCCTAC |  |
| 49 | L2HGDHsg2btm | AAACGTAGGGAATTCCCTTGAGGTC |  |
| 50 | L2HGDHsg3top | CACCGAAACATCCTGGACTTTCGAT |  |
| 51 | L2HGDHsg3btm | AAACATCGAAAGTCCAGGATGTTTC |  |
| 52 | L2HGDHsg4top | CACCGTCTTTTGATATAGTCATCGT |  |
| 53 | L2HGDHsg4btm | AAACACGATGACTATATCAAAAGAC |  |
| 54 | hKRT14_F | TGAGCCGCATTCTGAACGAG | Primers used for Real-Time qPCR analysis |
| 55 | hKRT14_R | GATGACTGCGATCCAGAGGA |  |
| 56 | hZEB1_F | CAGCTTGATACCTGTGAATGGG |  |
| 57 | hZEB1_R | TATCTGTGGTCGTGTGGGACT |  |
| 58 | mMyl1_F | AAGATCGAGTTCTCTAAGGAGCA |  |
| 59 | mMyl1_R | TCATGGGCAGAAACTGTTCAAA |  |
| 60 | mMyog_F | GAGACATCCCCCTATTTCTACCA |  |
| 61 | mMyog_R | GCTCAGTCCGCTCATAGCC |  |
| 62 | mMyl4_F | AAGAAACCCGAGCCTAAGAAGG |  |
| 63 | mMyl4_R | TGGGTCAAAGGCAGAGTCCT |  |
| 64 | mActc1_F | CTGGATTCTGGCGATGGTGTA |  |
| 65 | mActc1_R | CGGACAATTTCACGTTCAGCA |  |
| 66 | hACTC1_F | TCCCATCGAGCATGGTATCAT |  |
| 67 | hACTC1_R | GGTACGGCCAGAAGCATACA |  |
| 68 | hMYL1_F | GTTGAGGGTCTGCGTGTCTTT |  |
| 69 | hMYL1_R | ACCCAGGGTGGCTAGAAC |  |

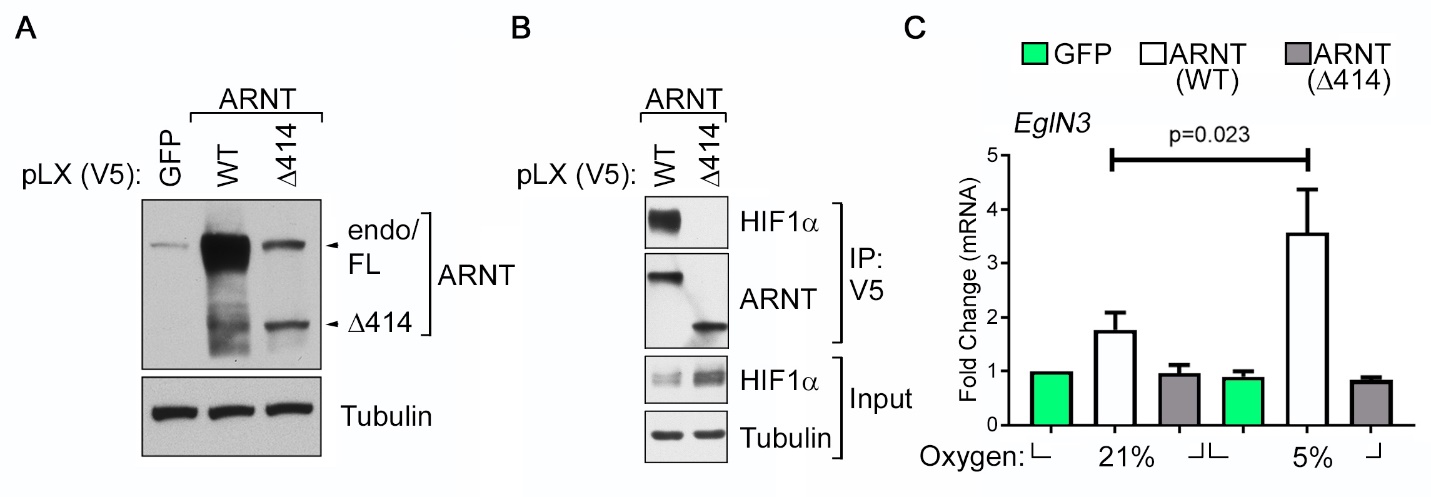

**Fig. S1. Re-Expression of Wild-Type ARNT Restores HIF Activity in *Arnt*-deficient mHepa-1 c4 Cells.**

(**A**) Anti-ARNT Immunoblot analysis of *Arnt*-deficient mHepa-1 c4 cells that were lentivirally transduced to produce V5-tagged indicated isoforms of ARNT or GFP (as a control) and cultured in 21% oxygen. FL = full-length. endo = endogenous. mHepa-1 c4 cells express a defective version of ARNT in which glycine 326 is replaced with aspartic acid (*32*). (**B**) Immunoblot analysis of anti-V5 immunoprecipitates from cell lysates of mHepa-1 c4 cells that express either wild-type or Δ414 ARNT cultured in the presence of 1 μM MLN for 24 hours. MLN was added to promote the accumulation of HIF1α. (**C**) Real-Time qPCR analysis of cells as in (**A**) that were cultured at the indicated oxygen concentration for 24 hours. Data represent mean±SD (n=3) and *p*-value was calculated using the Students *t*-test.

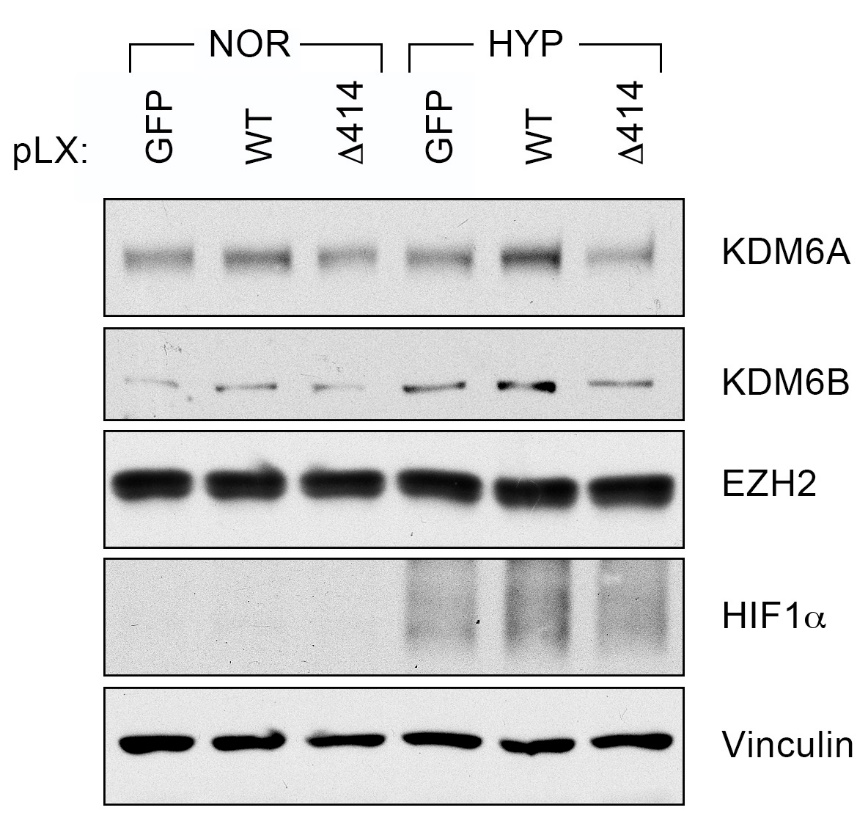

**Fig. S2. Expression Levels of H3K27 Modifiers in mHepa-1 c4 Cells**

Immunoblot analysis of mHepa-1 c4 cells that were lentivirally transduced to express the indicated ARNT isoforms or GFP (control) and cultured in 21% oxygen (NOR) or 5% oxygen (HYP), as indicated, for 2 days.

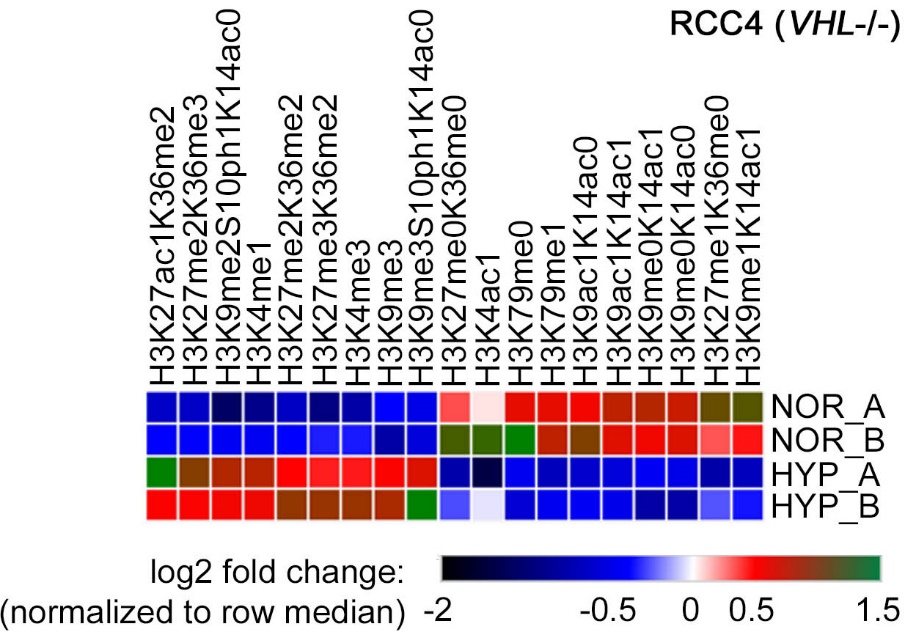

**Fig. S3. Hypoxia Induces Histone Hypermethylation in *VHL*-deficient Renal Cells.**

Histone modification profiling of 2 biological replicates of *VHL*-deficient RCC4 cells that were either cultured in 21% oxygen (NOR_A and NOR_B) or in 1% oxygen (HYP_A and HYP_B) for 3 days. The color in each cell represents the log_2_ fold change relative to all other samples in the column after first normalizing the total histone to an internal control peptide (Histone H3: residues 41-49).

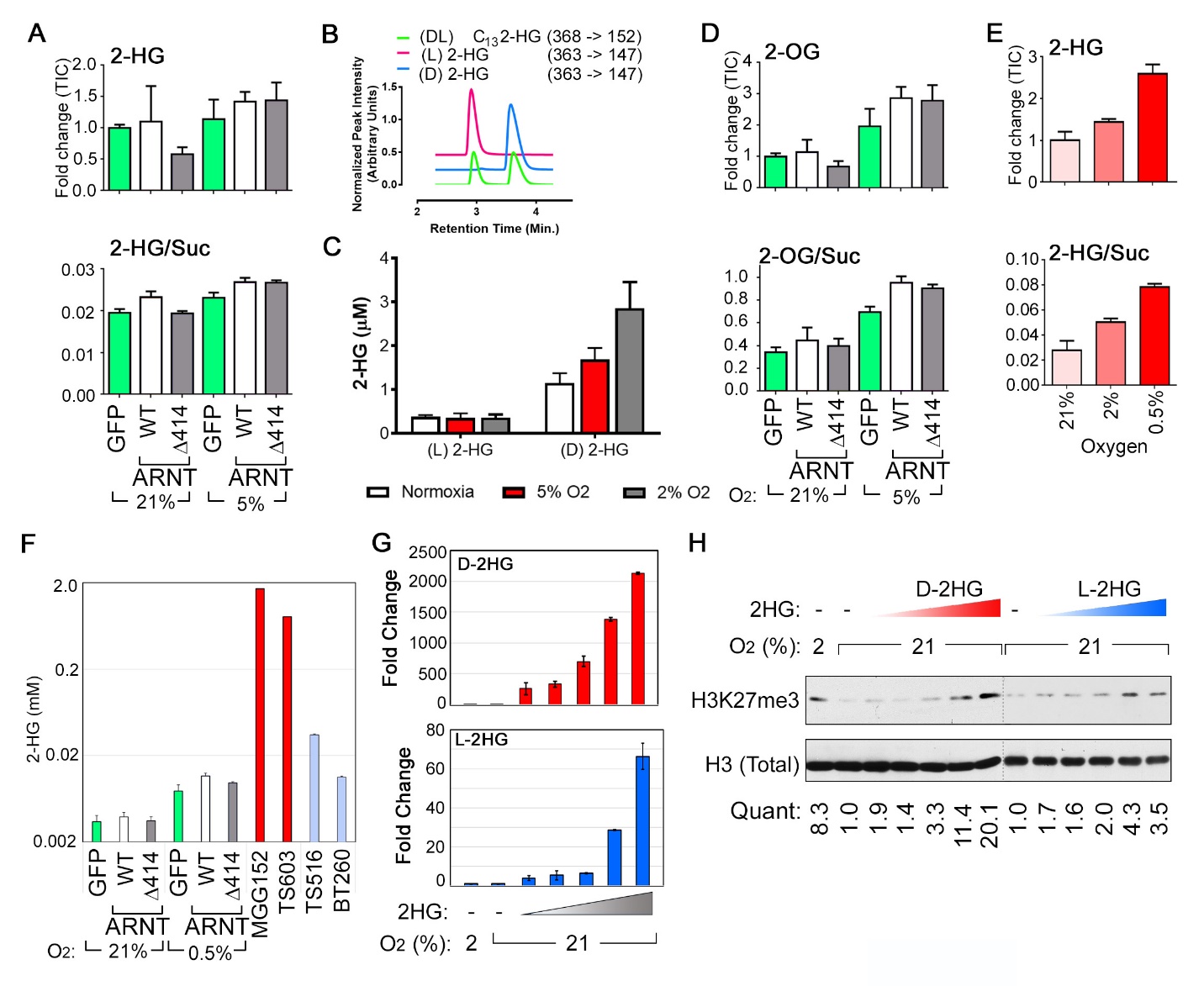

**Fig. S4. Increased Histone Methylation Observed After Exposure to Modest Hypoxia is Not Caused by 2-HG.**

(**A**) Intracellular levels of total 2-Hydroxyglutarate [2-HG] (**A**), as measured by GC-MS, in mHepa-1 c4 cells that were lentivirally transduced to produce the indicated ARNT isoforms or GFP (as a control) and cultured at the indicated oxygen concentrations for 4 days. Metabolite levels are represented either as fold changes in Total Ion Counts [TIC] [*top panel*] or as a ratio normalized to Succinate [Suc] levels [*bottom panel*]. Data represent mean±SD (n=4). (**B**-**C**) LC-MS analysis of a racemic mixture or individual enantiomers of 2-HG standards in mHepa-1 c4 cell matrix (**B**) and absolute intracellular 2-HG concentrations in parental Hepa-1 c4 cells grown at the indicated oxygen concentrations for 4 days (**C**). In (**C**), data represents mean±SD (n=3). (**D**) Intracellular levels of 2-oxoglutarate [2-OG] measured by GC-MS, as in (**A**). Data represents mean±SD (n=4). (**E**) Total 2-Hydroxyglutarate [2-HG], as measured by GC-MS, in *ARNT*-deficient mHepa-1 c4 cells that were cultured at the indicated oxygen concentrations for 4 days. Metabolite levels are represented either as fold changes in Total Ion Counts [TIC] [*top panel*] or as a ratio normalized to Succinate [Suc] levels [*bottom panel*]. Data represents mean±SD (n=4). (**F**) Absolute quantification of intracellular levels of 2-HG in the mHepa-1 c4 cells as in (**A**), cultured at the indicated oxygen concentrations for 4 days. IDH1 wild type [TS516 and BT260] or IDH1 mutant [MGG152 and TS603] glioma cell lines were included for comparison (*61*-*63*). Note use of log scale for y-axis. (**G**) Fold change in 2-HG relative to basal normoxic levels (set to 1) in parental mHepa-1 c4 cells that were cultured for 48 hours in the presence of increasing concentrations of esterified D-2HG (top panel) or L-2HG (bottom panel). Media was replaced every 24 hours and analysis was performed on cell lysates collected 3 hours after the last media change. Triangle represents a 3-fold serial dilution series ranging from 0, 19, 56, 167, 500, and 1500 µM concentration of the indicated enantiomer. Note that despite using identical concentrations for both enantiomers, the levels of intracellular D-2HG achieved was orders of magnitude higher than L-2HG, perhaps indicating differences in catabolic activity. Data represents mean±SD (n=3). (**H**) Immunoblot analysis of mHepa-1 c4 cells cultured at the indicated oxygen concentrations in the presence or absence of esterified 2-HG for 48 hours, as in (**G**). “Quant” represents fold-change in densitometric ratios of H3K27me3 (normalized to total H3) relative to untreated normoxic cells.

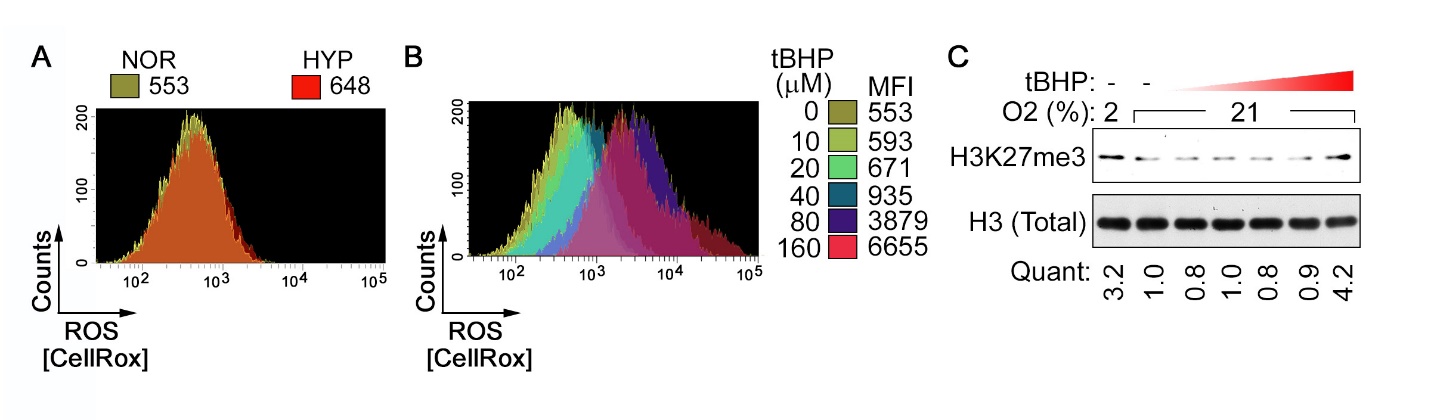

**Fig. S5. ROS Induction Does Not Account for the Effects of Hypoxia on Histone Hypermethylation.**

(**A** and **B**) Intracellular ROS levels, as measured by flow cytometry, in mHepa-1 c4 cells stained with the CellROX Deep Red reagent. (**A**) Comparison of cells cultured in 21% oxygen [NOR] or 2% oxygen [HYP], with values indicating Mean Fluorescence Index (MFI). (**B**) Comparison of intracellular ROS levels in cells treated with the indicated concentrations of tert-Butyl hydroperoxide (tBHP). The “tBHP=0” curve is identical to the “NOR” curve in (**A**) and is replicated for reference. (**C**) Immunoblot analysis of histones prepared from mHepa-1 c4 cells cultured at the indicated oxygen levels for 36 hours in the presence or absence of tBHP. Triangle represents a 2-fold serial dilution range of 10, 20, 40, 80, and 160 µM tBHP. “Quant” represents fold-change in densitometric ratios of H3K27me3 (normalized to total H3) relative to untreated normoxic cells.

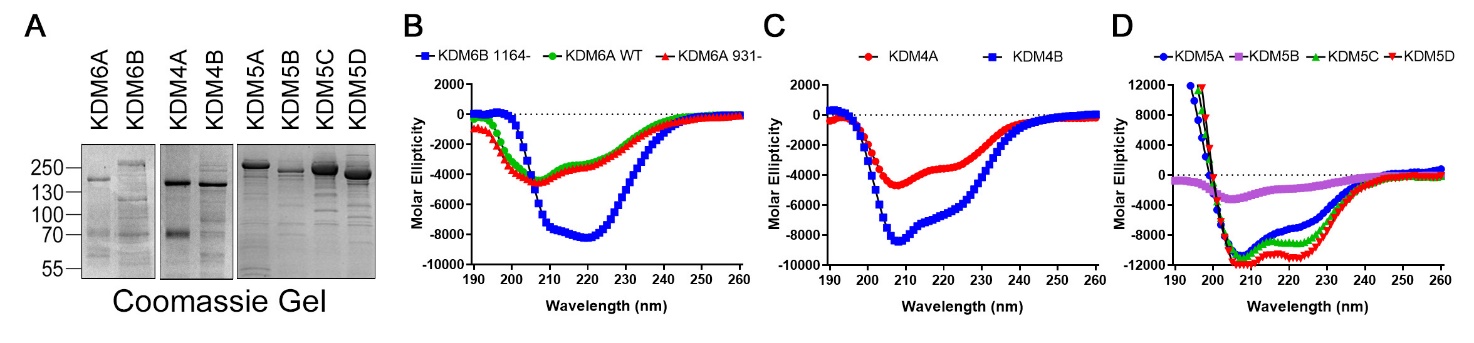

**Fig. S6. Purification of Recombinant Histone Demethylases.**

(**A**) Coomassie Blue dye staining of either His- or FLAG-tagged versions of the indicated histone demethylases that were expressed and affinity purified from baculovirally infected *Sf9* insect cells. “KDM6A 931-” = KDM6A*. “KDM6B 1164-“ = KDM6B*. (see text) (**B**) Circular Dichroism (CD) data that were collected between 190 and 260 nm at 22°C. Measurements were acquired every 1 nm with 1 s as an integration time and repeated three times with baseline correction.

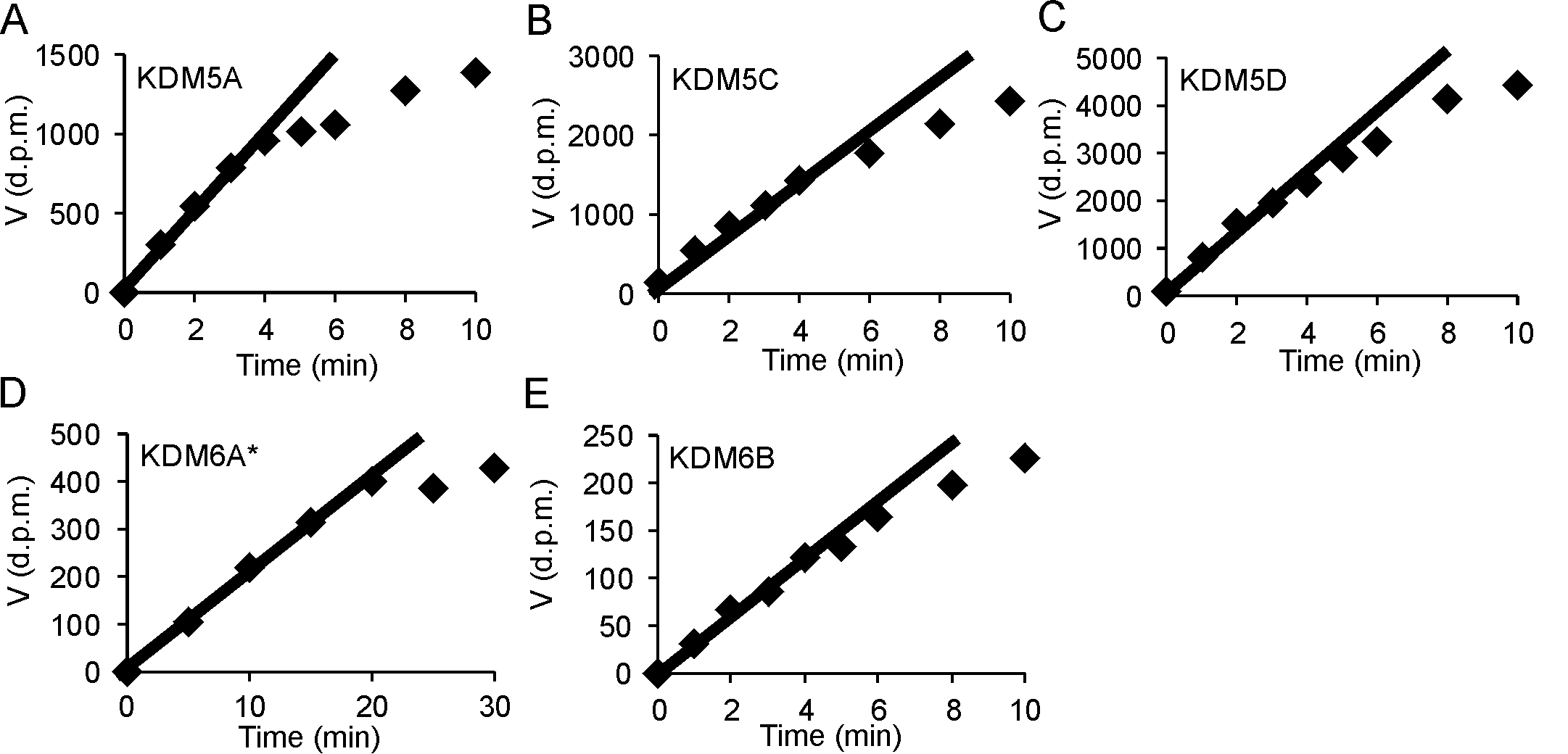

**Fig. S7. Time Course Analysis of Recombinant Purified Histone Demethylases.**

**(A**-**E**) The catalytic activity of the indicated recombinantly expressed and purified KDMs, as described in fig. S6, was studied with respect to time. The catalytic activity of KDM4A, KDM4B, KDM5B, KDM6A, and KDM6B* has been published earlier (*38*). * indicates the shortened version containing the JmjC domain.

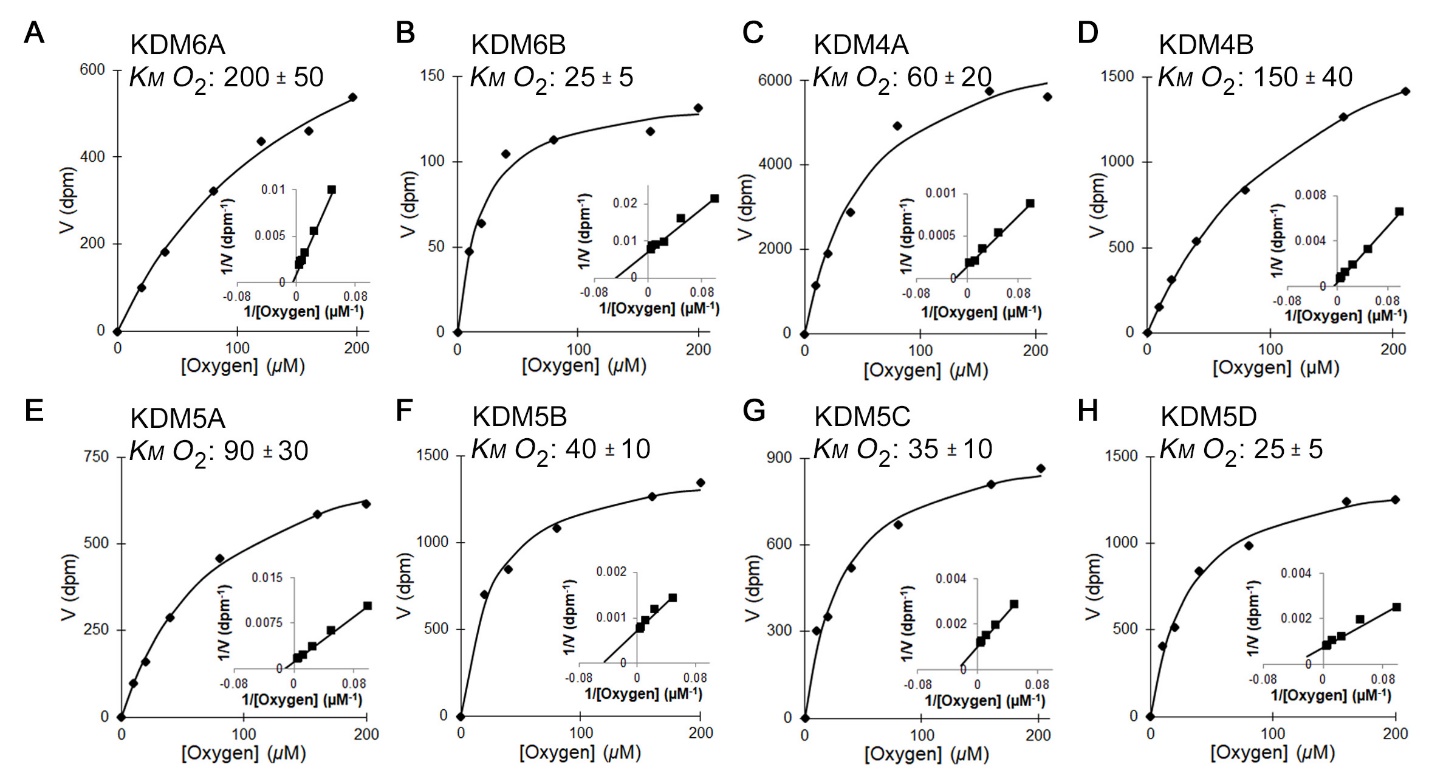

**Fig. S8. Biochemical Characterization of Recombinant Purified Histone Demethylases.**

(**A** – **H**) Representative Michaelis-Menten curves and Lineweaver-Burk plots [*insets*] showing the kinetic properties of the indicated baculovirally purified recombinant full length Histone Demethylases, measured over the indicated range of oxygen concentrations. *K_M_* values are mean±SD (n=3-12 independent measurements).

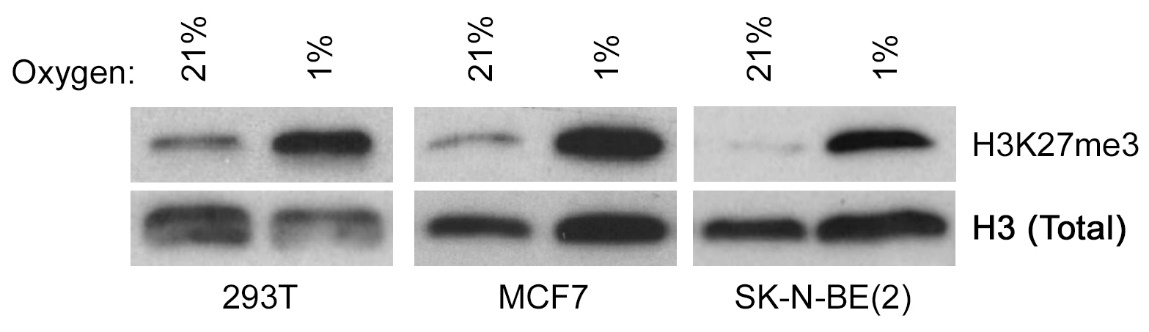

**Fig. S9. Hypoxia Induces Histone Hypermethylation in pVHL-proficient Cell Lines.**

Immunoblot analysis of histone lysates generated from the indicated cell lines cultured at the indicated oxygen concentrations for either 24 hours (MCF7 cells) or 72 hours (293T and SK-N-NE(2) cells).

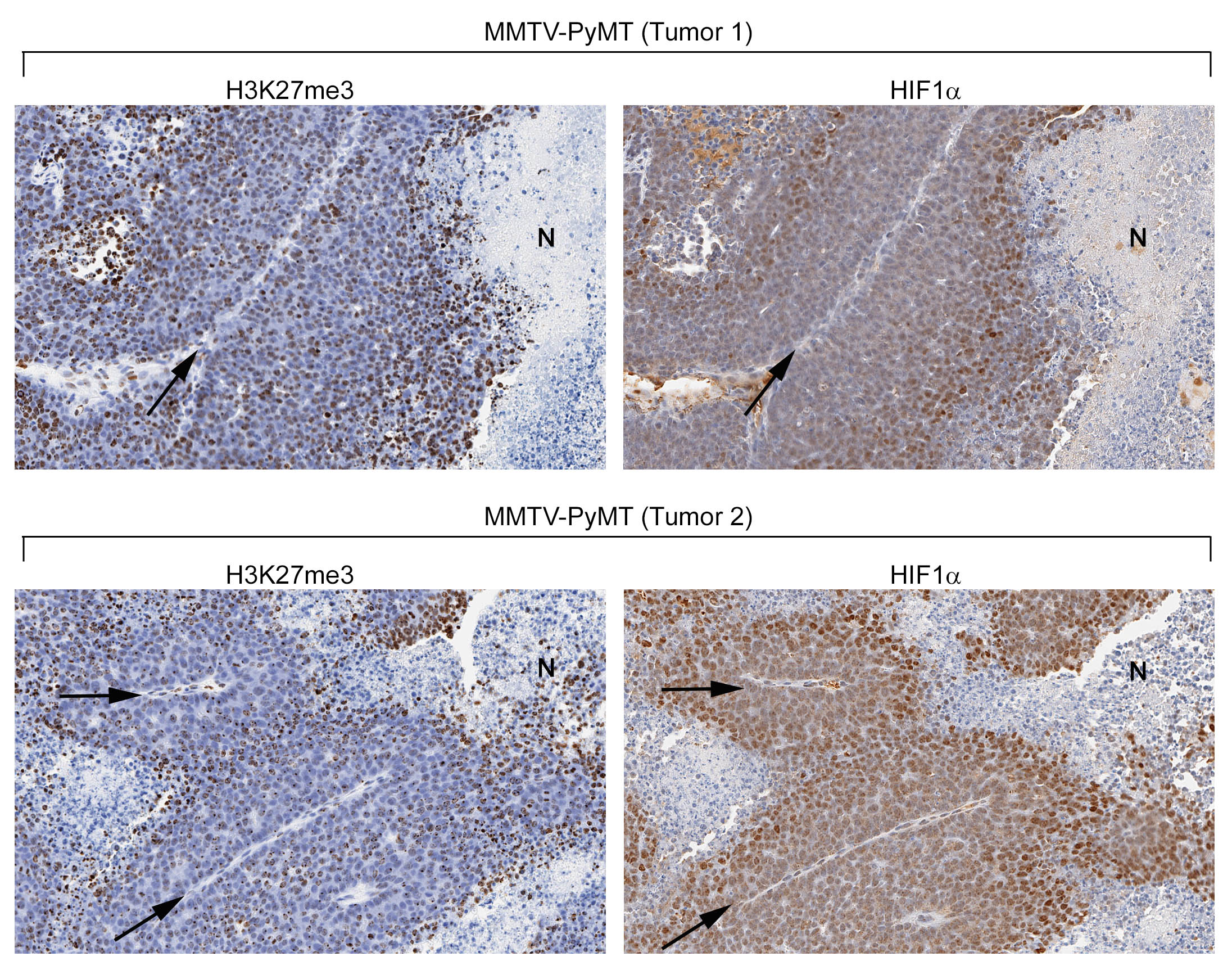

**Fig. S10. Tumor Hypoxia Promotes Histone Hypermethylation.**

Immunohistochemical analysis to compare localization of histone H3K27me3 and HIF1α, as indicated, in two independent breast tumors harvested from transgenic mice expressing mouse mammary tumor virus LTR-driven Polyoma virus middle T antigen [MMTV-PyMT. Blood vessel lumens are indicated by the arrows. Images are representative from analysis of 5 independent tumors.

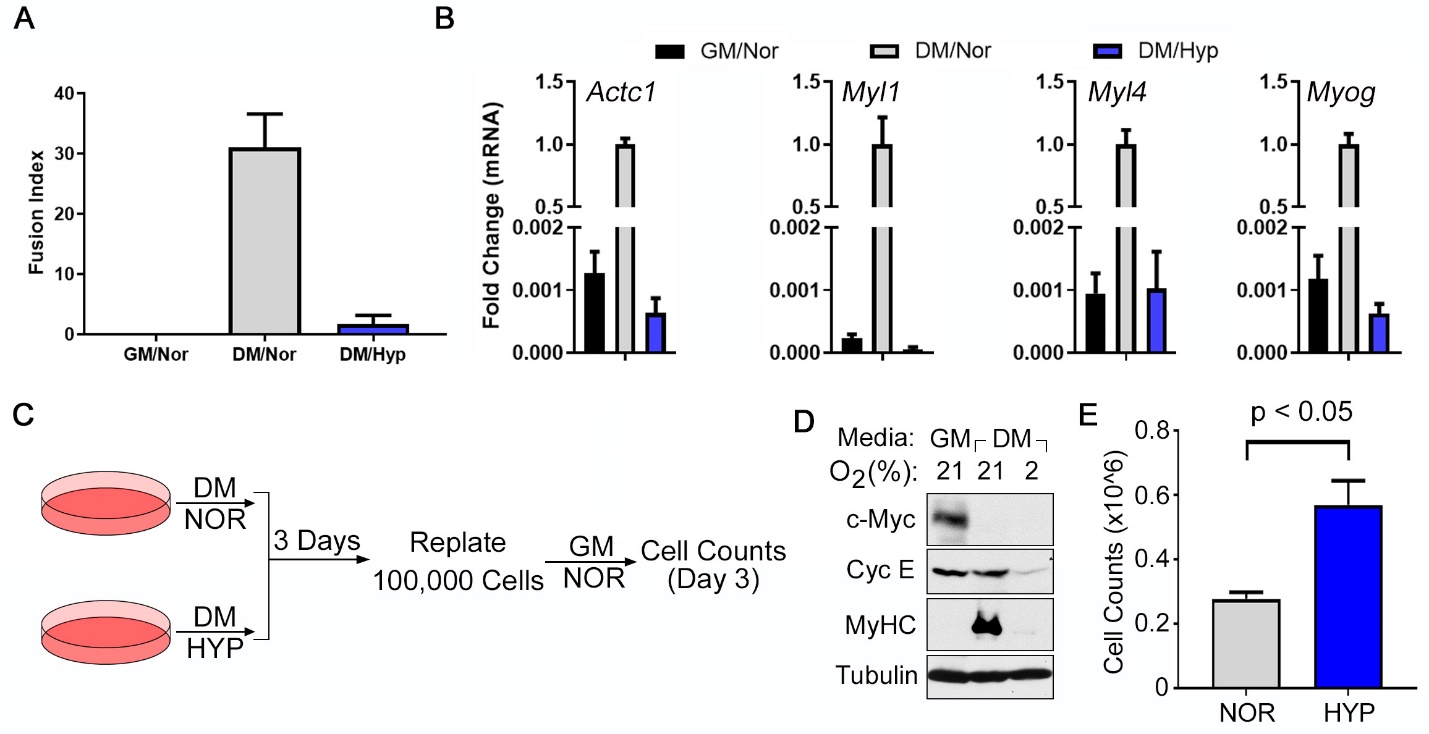

**Fig. S11. Hypoxia Blocks Differentiation by Arresting Cells in a Reversible Quiescence-like State**

**(Mathieu, #293)** Quantification of the effects of hypoxia either using fusion index measurements (**A**) or measuring mRNA levels of the indicated myogenic markers (**B**) in C2C12 cells that were cultured either at 21% (nor) or 2% oxygen (hyp) in the indicated media for 4 days. (**C**) Schema for testing the ability of C2C12 cells to proliferate after growth in DM under hypoxic conditions. (**D**) Immunoblot analysis of C2C12 cells after growth in DM under hypoxia or normoxia for 3 days, prior to replating as in (**C**). (**E**) Cell counts of viable C2C12 cells measured after replating in growth medium under normoxic conditions for 3 days as in (**D**). Data in (**A**), (**B**), and (**E**) represent mean±SD (n=3).

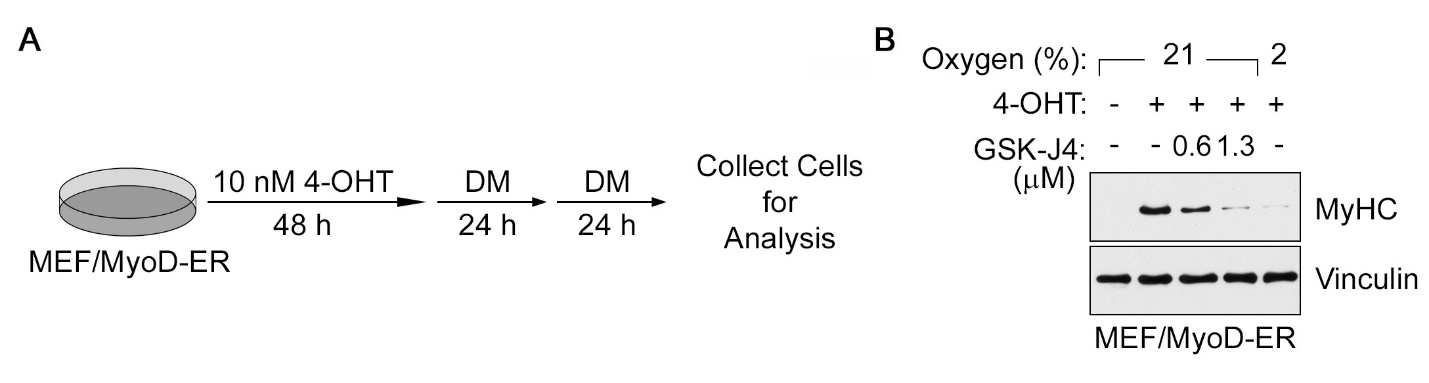

**Fig. S12. Hypoxia Blocks Myogenic Differentiation in MyoD-ER Expressing Mouse Embryonic Fibroblasts**

(Mathieu, #293) Experimental Scheme (**A**) and Immunoblot Analysis (**B**) of Mouse Embryonic Fibroblasts that were lentivirally transduced to express MyoD-ER, as previously described (*64*). In (**B**) cells were induced to differentiate as in (**A**) at the indicated oxygen concentrations in the presence or absence of GSK-J4. [4OHT = 4-Hydroxy Tamoxifen].

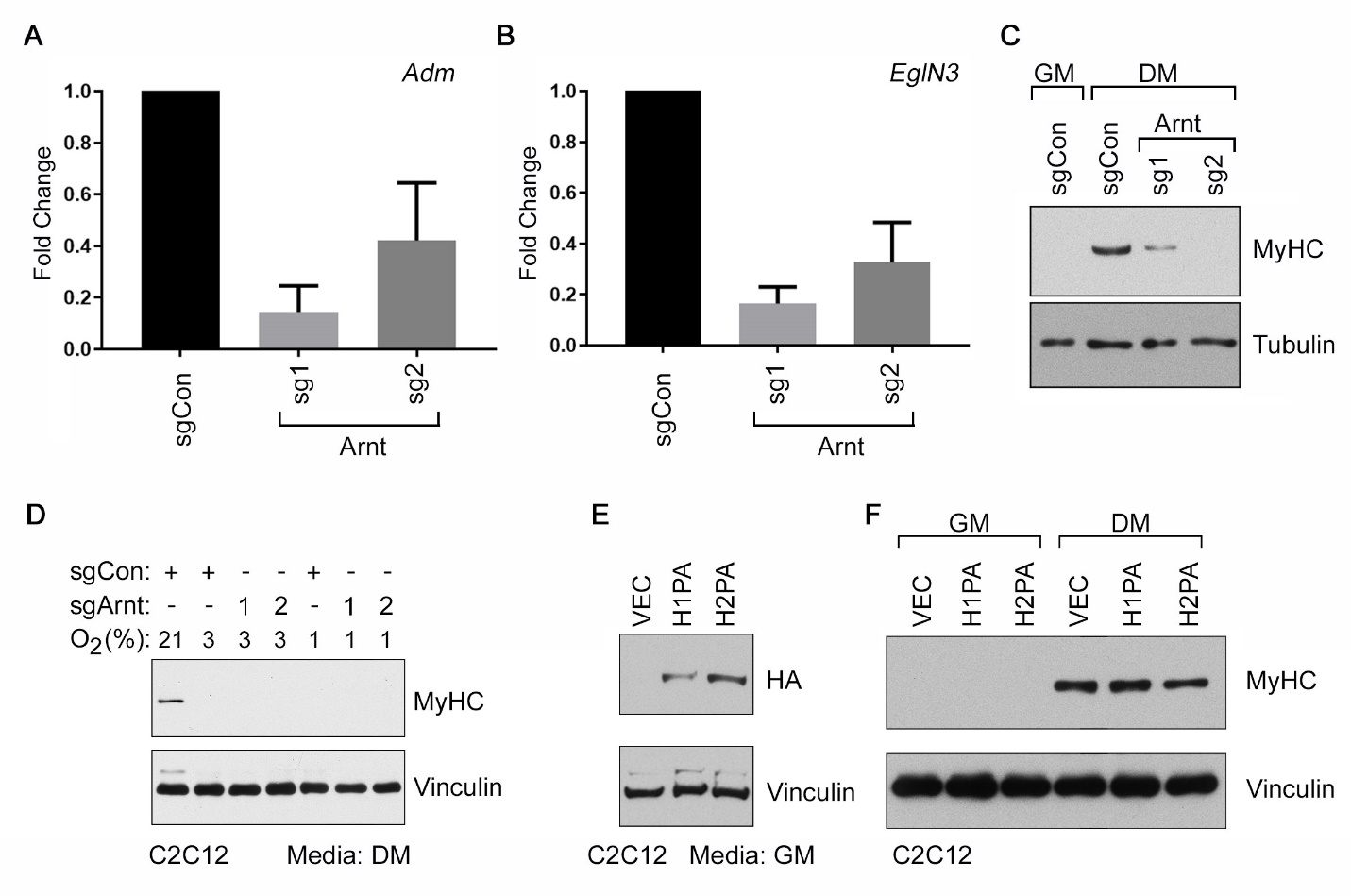

**Fig. S13. HIF Activity is Necessary for Differentiation and Not Sufficient to Block Differentiation of C2C12 Myoblasts.**

(**A**-**D**) Real-Time qPCR analysis of *Adm* (**A**) and *Egln3* (**B**) mRNAs and immunoblot analysis (**C** and **D**) of C2C12 cells that were lentivirally transduced to express the indicated sgRNAs targeting ARNT (ARNT sg1 and sg2) or a non-targeting control (sgCon) in the presence of Cas9. In (**A**) and (**B**) cells were cultured in growth medium at 2% oxygen for 24 hours and data represents mean±SD (n=3). In (**C**) cells were cultured either in growth medium [GM] or differentiation medium [DM], as indicated, in 21% oxygen for 4 days. In (**D**) cells were cultured in DM at the indicated oxygen concentrations for 4 days. (**E**-**F**) Immunoblot analysis of C2C12 cells that were lentivirally transduced to produced HA-tagged versions of mutant HIF1α [H1PA], mutant HIF2α [H2PA], or empty vector [VEC] as a control. In (**E**) cells were seeded in GM and harvested after 24 hours. In (**F**) cells were cultured in the indicated media in 21% oxygen for 4 days. The H1PA and H2PA variants contain alanine residues in place of the prolyl residues that are normally hydroxylated by the EglN prolyl hydroxylases (*49*).

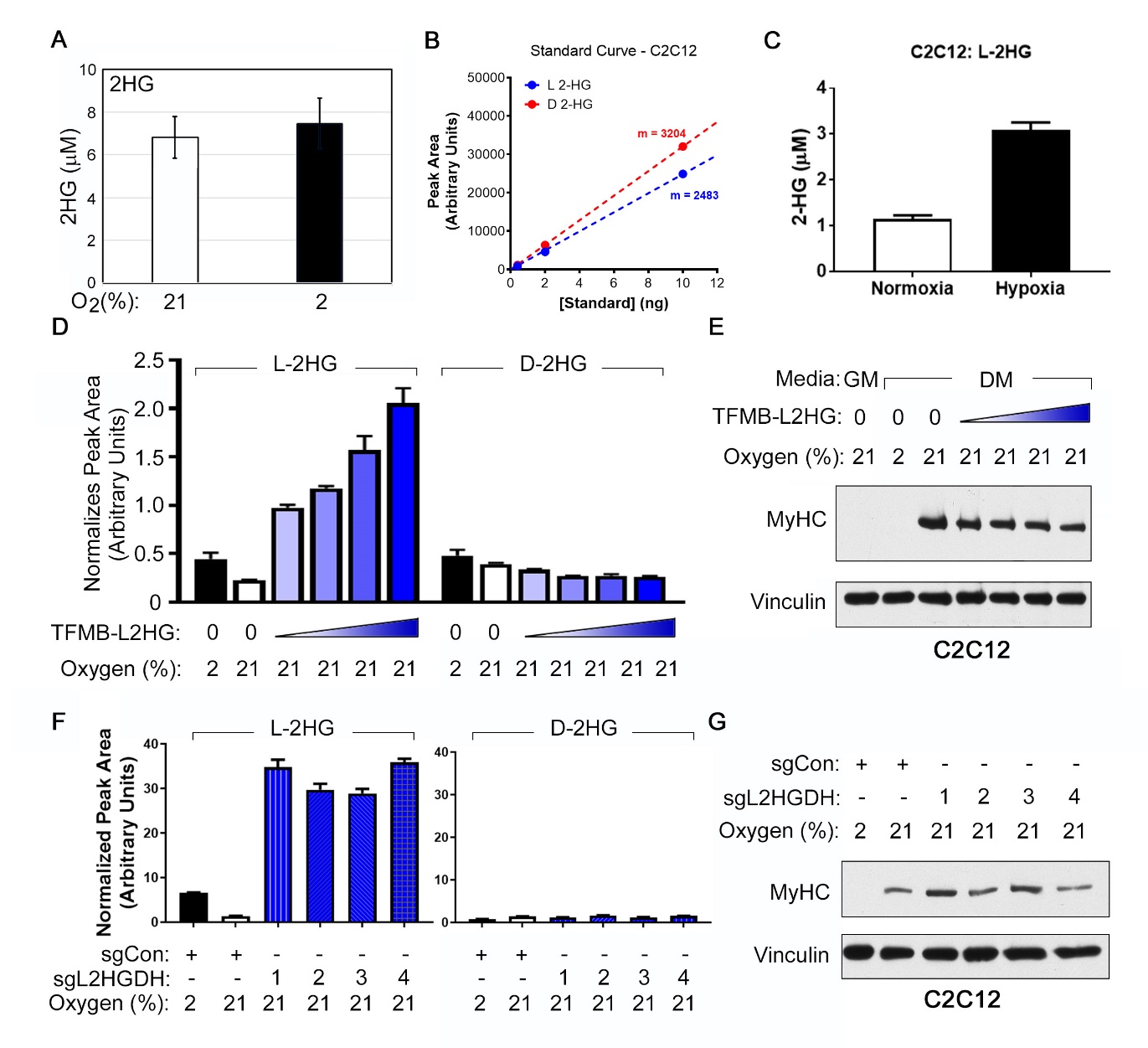

**Fig. S14. Hypoxic L-2HG Production is Not Sufficient to Block Differentiation in C2C12 Myoblasts.**

**(A)** Intracellular total 2HG levels in C2C12 cells determined by GC-MS, measured against a 2HG standard curve in water. (**B**-**C**) Intracellular L-2HG levels in C2C12 cells determined by LC-MS. Peak areas determined using a standard curve of ^13^C_5_-2-HG in C2C12 cell matrix (**B**) were used to measure absolute intracellular concentrations of L-2HG (**C**) in C2C12 cells that were cultured for 4 days in differentiation media at either 21% oxygen (Normoxia) or 2% oxygen (Hypoxia). (**D)** Intracellular enantiomer-specific 2-HG levels, normalized to the spiked-in ^13^C_5_-2-HG internal standard, as determined by LC-MS. Triangles indicate treatment with a 3-fold serial dilution range (19, 56, 167, and 500 µM) of an esterified (TFMB) version of L-HG**.** Esterified L-2HG was replenished at every daily media change and measurements were made 3 hours after the last media change. (**E**) Immunoblot analysis of C2C12 cells treated with TFMB-L2HG as in (**D**) in the indicated media and oxygen environments. (**F**) Intracellular enantiomer-specific 2-HG levels, normalized to the spiked-in ^13^C_5_-2-HG internal standard, as determined by LC-MS, in C2C12 cells that were lentivirally transduced to express the indicated L2HGDH (sgL2HGDH) sgRNAs or control (sgCon) and cultured at the indicated oxygen concentrations in differentiation media for 4 days. (**G**). Immunoblot analysis of C2C12 cells as in (**F**) grown in DM at the indicated oxygen concentrations. Data in (**A**), (**C**), (**D**), and (**F**) represent mean±SD (n=3).

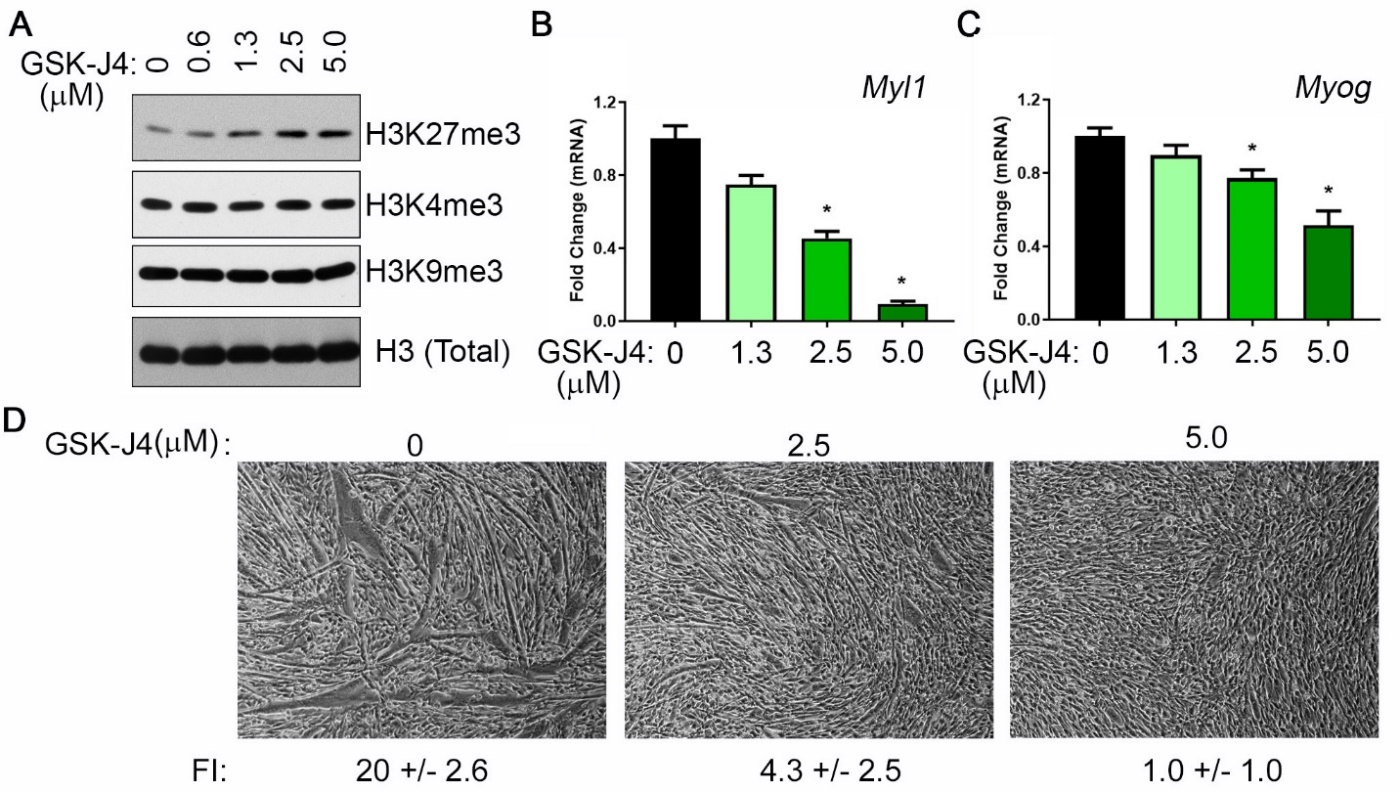

**Fig. S15. Treatment with GSK-J4 Blocks Differentiation in C2C12 Myoblasts.**

(**A**) Immunoblot analysis of histone lysates generated from C2C12 cells treated with the indicated concentrations of GSK-J4 for 3 days. (**B**-**D**) Fold change in mRNA levels (normalized to *Actin*) of *Myl1* (**B**) and *Myog* (**C**), as measured by RealTime-qPCR, and Photomicrographs (**D**) of C2C12 cells that were cultured in differentiation media in the presence of the indicated concentrations of GSK-J4 for 4 days. In (**B**) and (**C**), data represents mean±SD (n=3) and [*] represents p<0.05 calculated by Students *t-test*. In (**D**), Fusion Index (FI) is represented as mean±SD (n=3).

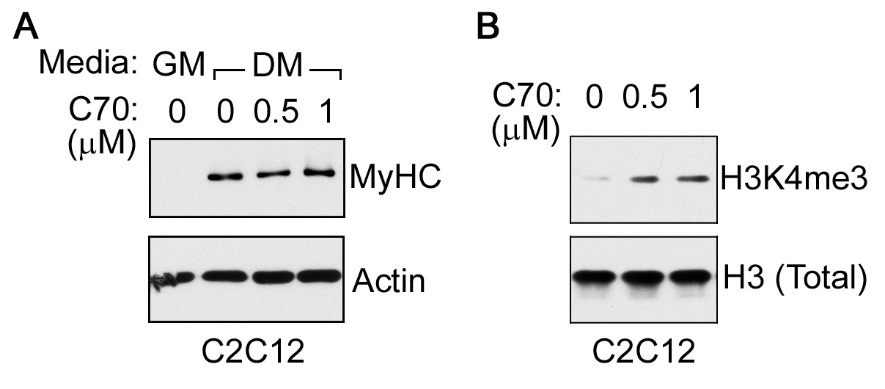

**Fig. S16. Treatment with KDM-C70 Does Not Block the Differentiation of C2C12 Myoblasts.**

(Mathieu, #293) Immunoblot analysis of soluble protein (**A**) and histone lysates (**B**) generated from C2C12 cells cultured in the indicated media in (**A**) or in DM in (**B**) in the presence of the indicated concentrations of KDM-C70 for 4 days.

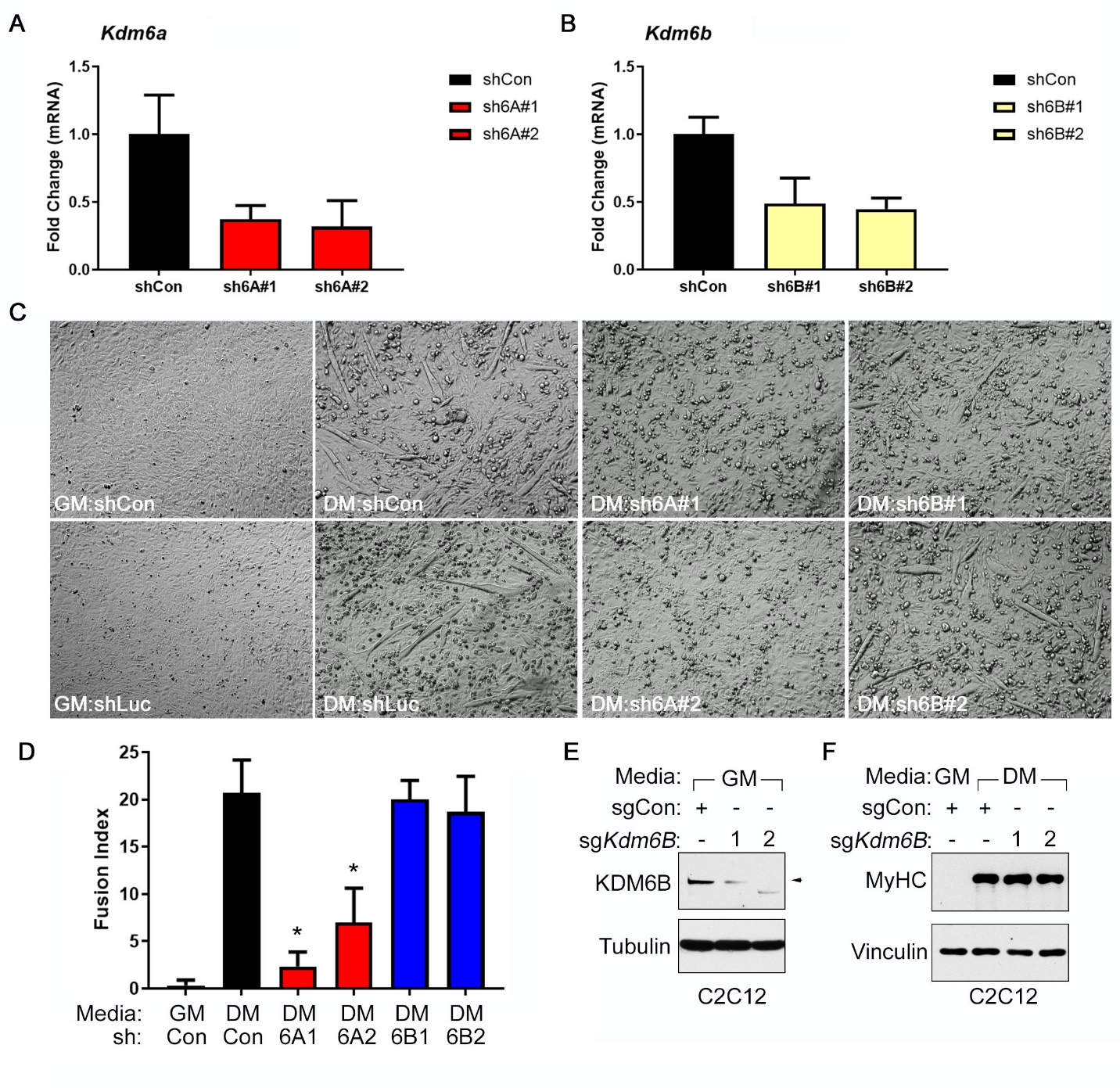

**Fig. S17. KDM6B Inactivation Does Not Block Differentiation in C2C12 Myoblasts.**

(Mathieu, #293) Real-Time qPCR analysis to measure *Kdm6A* (**A**) and *Kdm6B* (**B**) mRNAs in C2C12 cells that were lentivirally transduced to express short-hairpin RNAs targeting *Kdm6A* [sh6A#1 and sh6A#2], *Kdm6B* [sh6B#1 and sh6B#2], or a non-targeting control shRNA [shCon]. Data represent mean±SD (n=3). (**C**-**D**) Photomicrographs (**C**) and Fusion Index Measurements (**D**) of C2C12 cells that were lentivirally transduced to express the indicated shRNAs and then cultured in the indicated media for 4 days. In (**D**), data represent mean±SD (n=3), and [*] indicates p<0.05 calculated by Students *t-test*. (**E**-**F**) Immunoblot analysis from C2C12 cells lentivirally transduced to express *Kdm6B* sgRNA or control (Con) sgRNA. In (**E**) cells cultured in the indicated media were analyzed 7 days post infection. In (**F**) cells were cultured in the indicated media for 4 days.

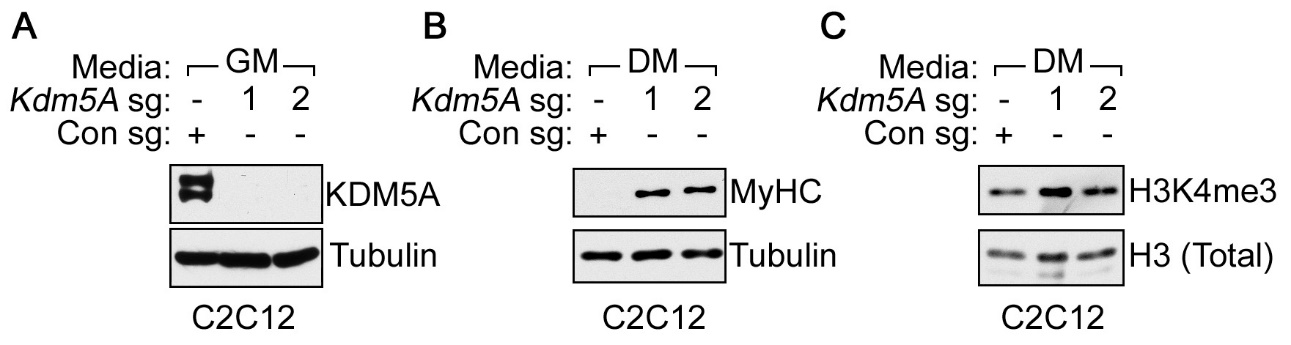

**Fig. S18. KDM5A Inactivation Promotes Myogenic Differentiation in C2C12 Cells.**

(**A**-**C**) Immunoblot analysis of soluble proteins (**A** and **B**) and histone lysates (**C**) from C2C12 cells that were lentivirally transduced to express the indicated *Kdm5A* sgRNAs or control (Con) sgRNA. In (**A**) cells cultured in the indicated media were analyzed 7 days post infection. In (**B**) and (**C**) cells were cultured in the indicated media for 4 days. Note that in panel (**B**) the absence of a MyHC signal in lane 1 is because it was necessary to do a very short exposure due to the strong MyHC signal in lanes 2 and 3.

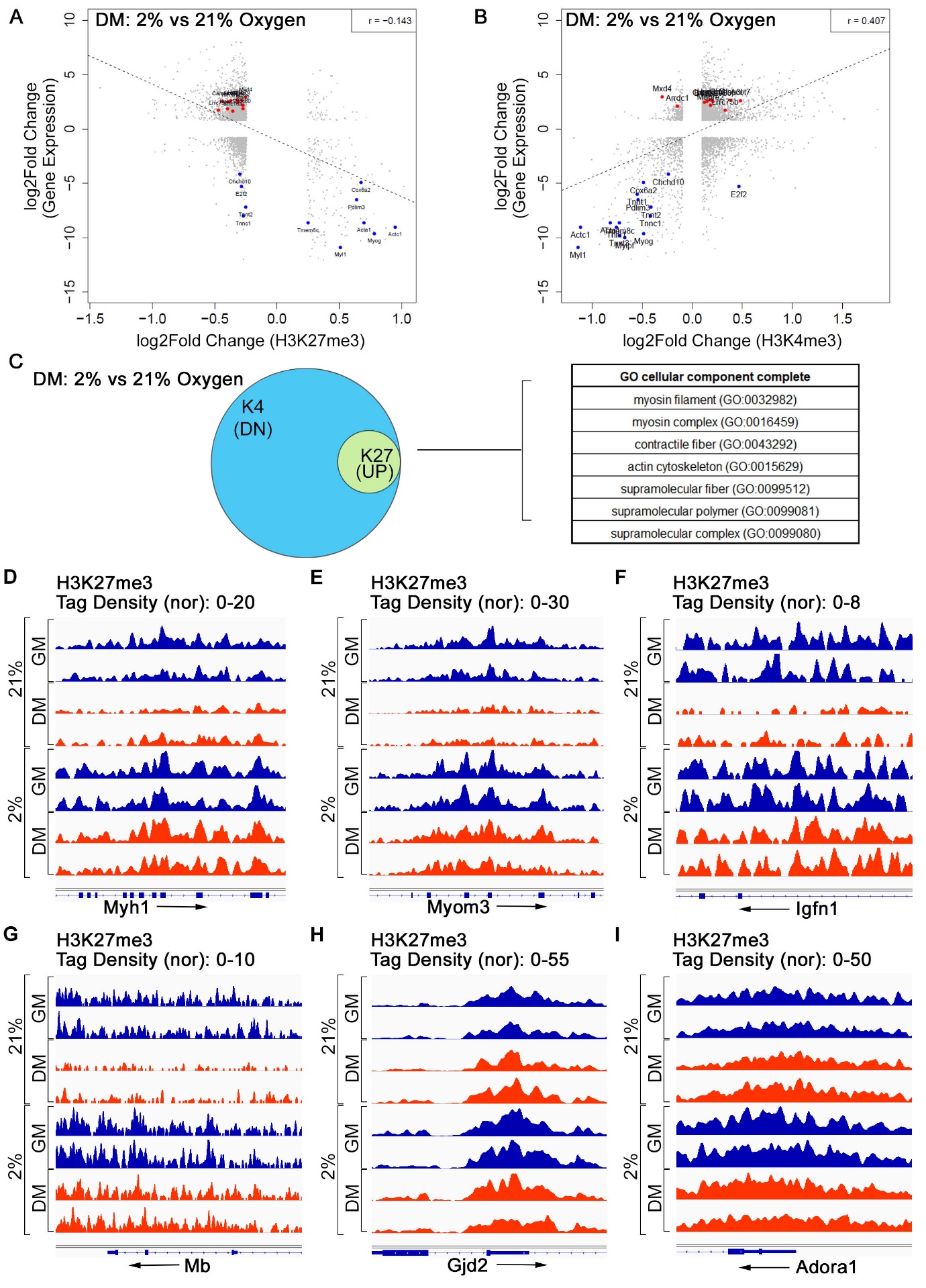

**Fig. S19. Genomic Analysis Demonstrates Failure to Erase H3K27me3 at Myogenic Regulators in Hypoxic C2C12 Cells.**

(**A**-**C**) Correlation of genomic recruitment (as measured by ChIP-Seq) versus gene expression (as measured by RNA-Seq) for H3K27me3 (**A**) and H3K4me3 (**B**) in C2C12 cells that were cultured in differentiation media (DM) at either 21% or 2% oxygen. Gene Ontology Enrichment in (**C**) comparing biological processes that show increased recruitment of H3K27me3 and decreased recruitment of H3K4me3 in hypoxia (2%) versus normoxic (21%) cells. (**D**-**I**) H3K27me3 recruitment, as measured by tag density from ChIP-Seq data, in C2C12 cultured under the indicated conditions for 4 days. Recruitment at the myogenic markers *Myh1* (**D**), *Myom3* (**E**), *Igfn1* (**F**), and *Mb* (**G**) [and *Actc1*, *Myl1*, and *Myog* shown in Fig. 4, **D** to **F**]. *Gjd2*, which lies adjacent to *Actc1*, and *Adora1*, which lies adjacent to *Myog*, were included as specificity controls.

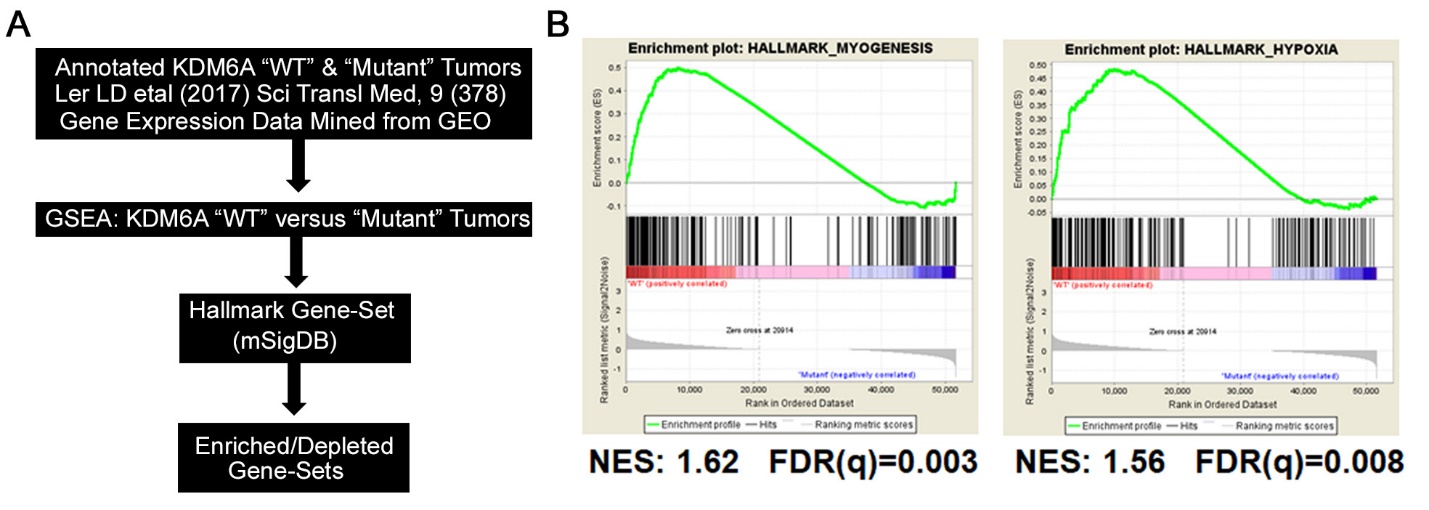

**Fig. S20. KDM6A Loss is Associated with Hallmarks of Dedifferentiation in Bladder Tumors**

(Mathieu, #293) Analytical flowchart (**A**) and Gene Set Enrichment Analysis showing the enrichment plots for transcriptional signatures that are enriched in KDM6A wild-type “WT” versus KDM6A “Mutant” tumors (n=10 in each group). Note that the wild-type KDM6A tumors have increased expression of “Hallmark Myogenesis” mRNAs relative to KDM6A mutant tumors. This does not appear to be because the KDM6A mutant tumors are more hypoxic if one examines levels of “Hallmark Hypoxia” mRNAs, which are actually higher in the wild-type KDM6A tumors. FDR (q) values below 0.25 were considered statistically significant.

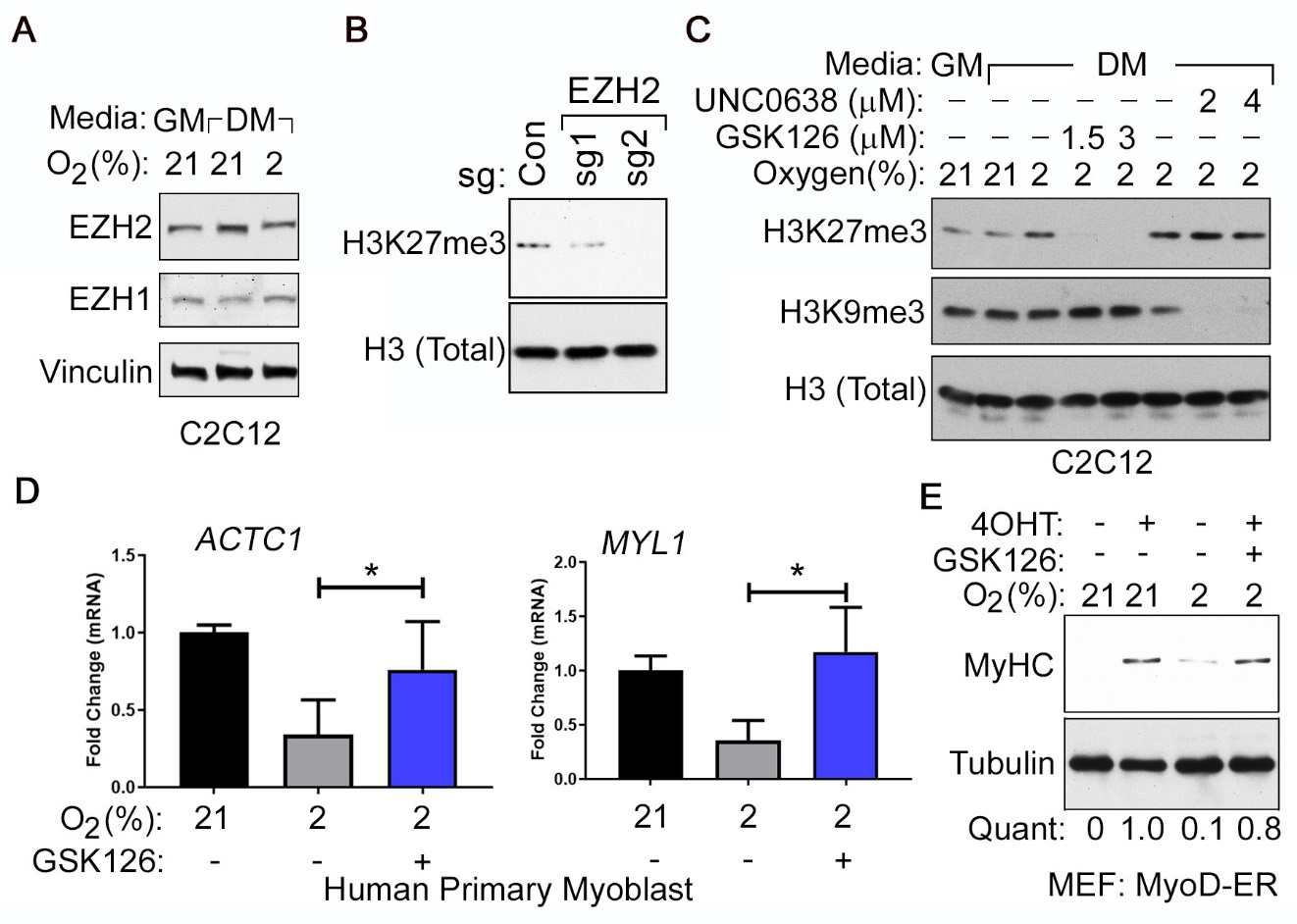

**Fig. S21. Pharmacological Inhibition of EZH2 Rescues the Hypoxic Differentiation Block in Myogenic Cells**

(**A)** Immunoblot analysis of C2C12 myoblasts cultured in the indicated media and oxygen concentrations for 2 days. (**B**) Immunoblot analysis of histone lysates prepared from C2C12 myoblasts that were transduced to express either *Ezh2* sgRNAs or a control (Con) sgRNA, as indicated. Histones were extracted 7 days post infection. (**C**) Immunoblot analysis of histone lysates generated from C2C12 cells treated with the indicated concentrations of GSK126 or UNC0638 for 4 days. Cells were cultured either in 21% oxygen (NOR) or 2% oxygen (HYP) in growth medium (GM) or differentiation medium (DM), as indicated. (**D**) mRNA levels of myogenic differentiation markers, relative to Actin mRNA, measured by Real-Time qPCR of RNA obtained from primary human skeletal myoblasts that were cultured in differentiation media at the indicated oxygen concentrations in the presence or absence of 2 µM GSK126, as indicated. Data represent mean±SD (n=3), and [*] indicates p<0.05 calculated by Students *t-test*. (**E**) Immunoblot analysis of MyoD-ER expressing Mouse Embryonic Fibroblasts (described in fig. S12) cultured in differentiation media under the indicated oxygen concentrations in the presence or absence of 2 µM GSK126, as indicated. “Quant” represents fold-change in densitometric ratios of MyHC (normalized to Tubulin) relative to normoxic cells treated with 4-OHT. [4OHT=4-Hydroxy Tamoxifen].

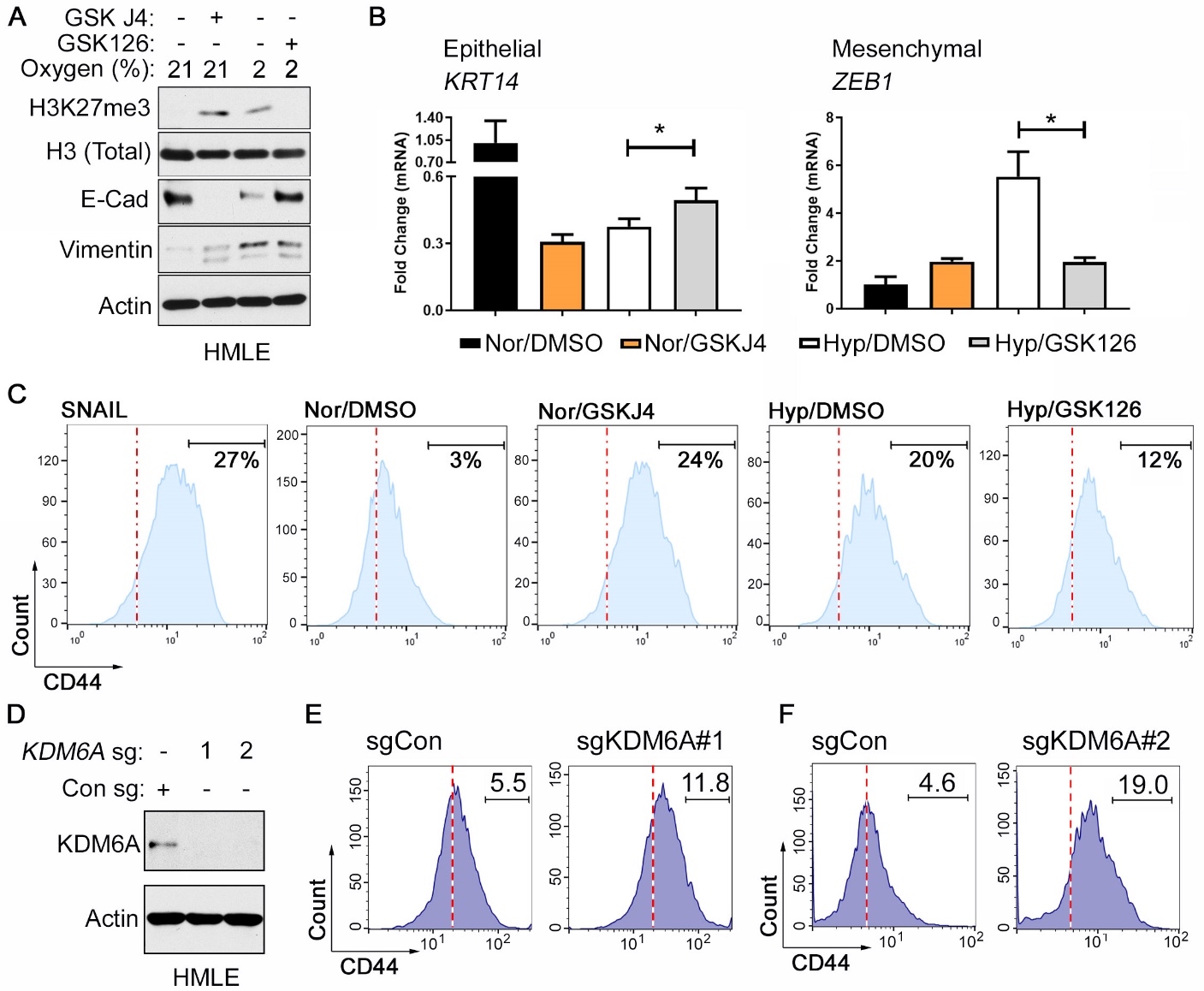

**Fig. S22. Modulating H3K27me3 Regulates EMT in Human Mammary Epithelial (HMLE) Cells**

(**A**-**C**) Immunoblot Analysis (**A**), mRNA levels of EMT markers (relative to Actin) determined by Real-time qPCR analysis (**B**), and flow cytometric analysis (**C**), of HMLE cells that were cultured at 21% Oxygen (Nor) or 2% Oxygen (Hyp) for 7 days in the presence 2 µM GSK-J4, 3 µM GSK126, or DMSO as indicated. In (C), SNAIL represents data from HMLE cells lentivirally transduced to express the transcription factor SNAIL, a master regulator of EMT. (**D**-**F**) Immunoblot analysis (**D**) and flow cytometric analysis (**E** and **F**) of HMLE cells that were lentivirally transduced to express one of two different *KDM6A* sgRNAs or a control sgRNA and cultured under normoxic conditions for 7 days post selection. Data in (**B**) represent mean±SD (n=3). (**C**), (**E**), and (**F**) is representative of 3 independent experiments.

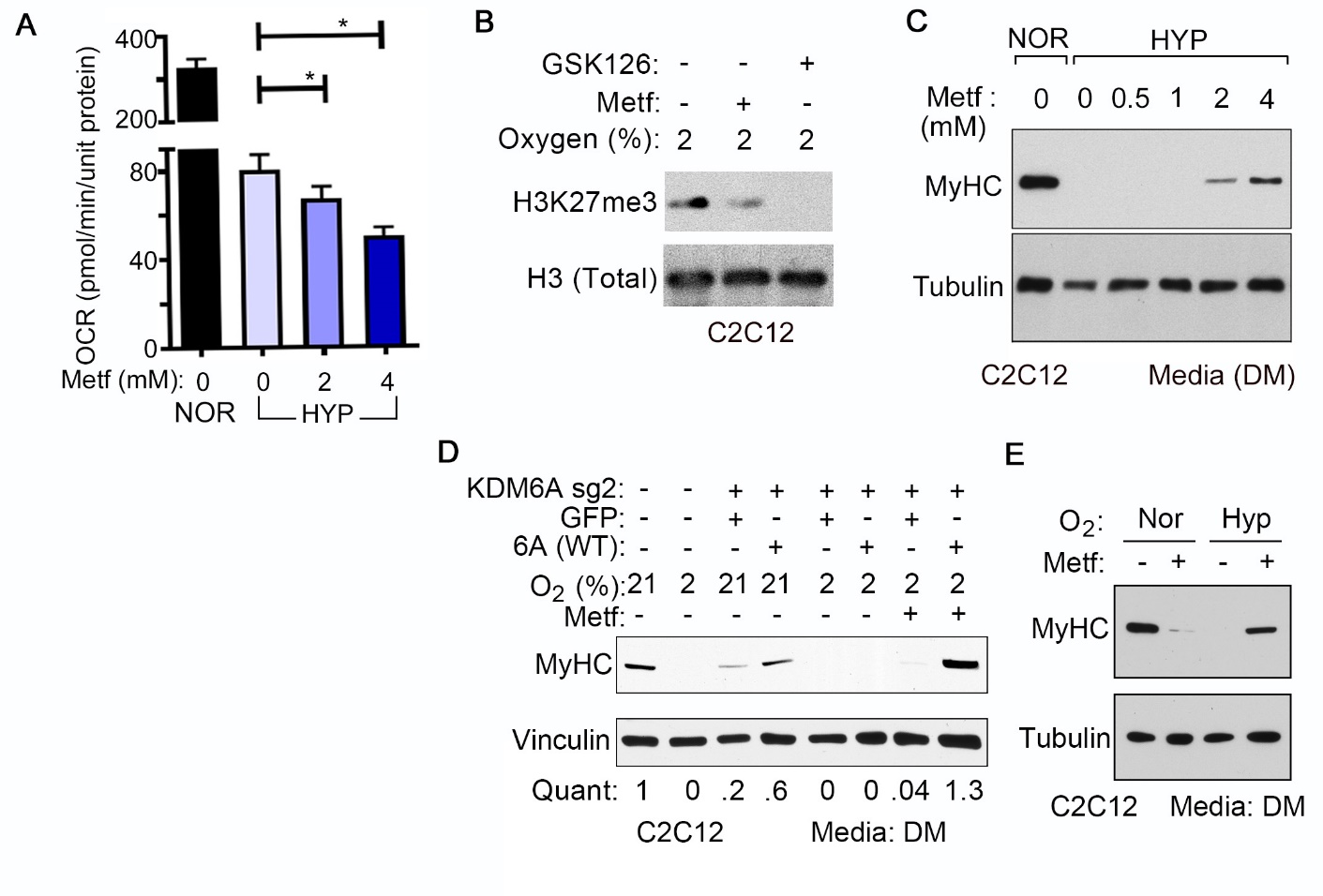

**Fig. S23. Metformin Treatment Decreases Oxygen Consumption in Hypoxic Cells and Rescues Differentiation in a KDM6A-dependent Manner.**

(**A**) Basal Oxygen Consumption Rate [OCR] as determined by Seahorse measurements in C2C12 cells that were cultured in 21% oxygen [NOR] or 2% oxygen [HYP] and treated with the indicated concentrations of Metformin [Metf]. (**B**) Immunoblot analysis of histone lysates generated from C2C12 cultured for 4 days in 2% oxygen in the presence of 2 mM Metformin [Metf], 3 µM GSK126, or DMSO [Unt=Untreated], as indicated. (**C**) Immunoblot analysis of C2C12 cells that were cultured in differentiation medium in 21% oxygen [NOR] or 2% oxygen [HYP] in the presence of the indicated concentrations of Metformin [Metf]. (**D**) Immunoblot analysis of C2C12 cells expressing, where indicated, *Kdm6a* sg2 [described in Fig. 3, (**F**) and (**G**)] that were lentivirally transduced to produce either wild-type human KDM6A (WT) or GFP and cultured in Differentiation Medium at the indicated oxygen concentrations for 4 days either in the presence or absence of 2mM Metformin (Metf), as indicated. “Quant” represents fold-change in densitometric ratios of MyHC (normalized to Vinculin) relative to untreated parental cells cultured under normoxic conditions (lane 1). (**E**) C2C12 cells cultured in Nor [=21% oxygen] or Hyp [=2% oxygen], as indicated. In both (**D**) and (**E)** cells were cultured in the presence of 2 mM Metformin [Metf] where indicated.

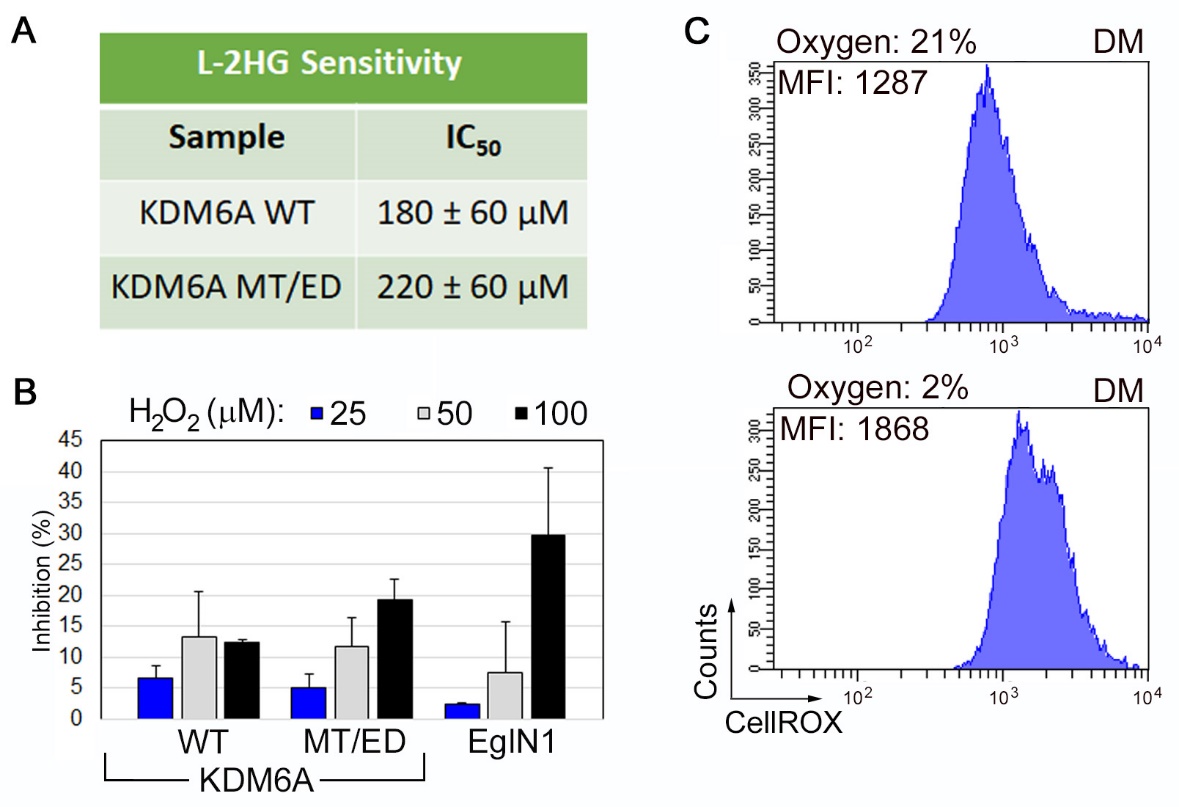

**Fig. S24. Wild-Type and Mutant KDM6A are Equally Insensitive to L-2HG and ROS**

**(A)** IC_50_ values of L-2HG as determined in vitro using recombinant wild-type KDM6A or the MT/ED mutant. (**B**) Percent inhibition in activity of the indicated proteins upon exogenous addition of the indicated concentrations of H_2_O_2_ (25 or 50 μM). Inhibition was calculated as a percentage loss in activity compared to reactions not containing H_2_O_2_. (**C**) Intracellular ROS levels as measured by flow cytometry after CellROX staining in C2C12 cells that were cultured in differentiation media (DM) at the indicated oxygen concentrations for 3 days. In (**A**) and (**B**), data represent mean±SD (n=3). In (**C**), data is representative of 3 independent measurements. [MFI = Mean Fluorescence Index].

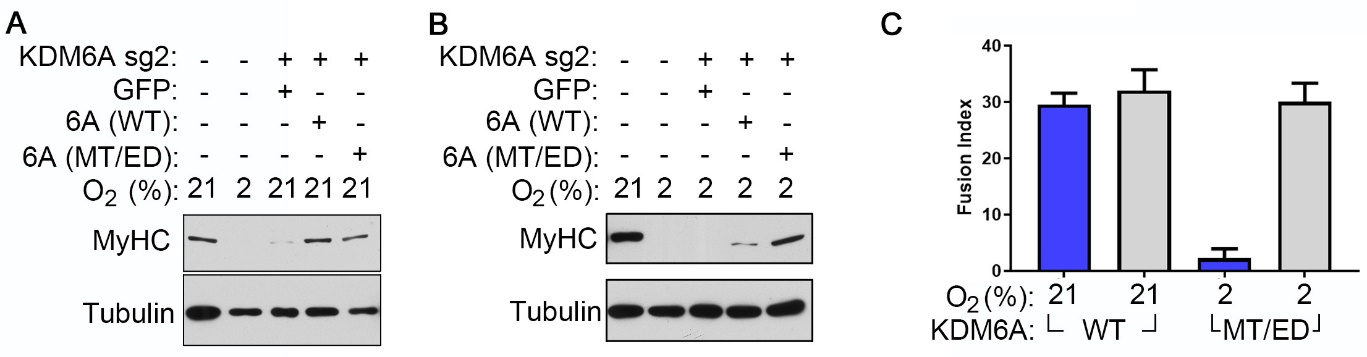

**Fig. S25. Reintroduction of an Enhanced Oxygen Affinity KDM6A Mutant is Sufficient to Rescue the Hypoxic Differentiation Block in C2C12 Cells**

(Mathieu, #293) Immunoblot analysis of C2C12 cells expressing, where indicated, *Kdm6a* sg2 [described in Fig. 3, (**F**) and (**G**)] that were lentivirally transduced to produce wild-type human KDM6A [6A(WT)], the human KDM6A MT/ED double mutant [6A(MT/ED)], or GFP (control), and then cultured in DM at the indicated oxygen concentrations for 4 days. (**C**) Fusion Index measurements for cells described in Fig. 5M. Data represent mean±SD (n=3).
